# Metabolic modeling of *Pectobacterium parmentieri* SCC3193 provides insights into metabolic pathways of plant pathogenic bacteria

**DOI:** 10.1101/284968

**Authors:** Sabina Zoledowska, Luana Presta, Marco Fondi, Francesca Decorosi, Luciana Giovannetti, Alessio Mengoni, Ewa Lojkowska

## Abstract

Understanding the plant-microbe interactions are crucial for improving plant productivity and plant protection. The latter aspect is particularly relevant for sustainable agriculture and development of new preventive strategies against the spread of plant diseases. Constraint-based metabolic modeling is providing one of the possible ways to investigate the adaptation to different ecological niches and may give insights into the metabolic versatility of plant pathogenic bacteria. In this study, we present a curated metabolic model of the emerging plant pathogenic bacterium *Pectobacterium parmentieri* SCC3193. Using flux balance analysis (FBA), we predict the metabolic adaptation to two different ecological niches, relevant for the persistence and the plant colonization by this bacterium: soil and rhizosphere. We performed *in silico* gene deletions to predict the set of core essential genes for this bacterium to grow in such environments. We anticipate that our metabolic model will be a valuable element for defining a set of metabolic targets to control infection and spreading of this plant pathogen and a scaffold to interpret future –omics datasets for this bacterium.

## INTRODUCTION

Plant-bacteria interplays have been studied over long periods of time, mainly in terms of pathogenic and beneficial (symbiotic) interactions. Various details are now known concerning the molecular basis of all these interactions^1^. For example, biological studies of plant pathogenic bacteria allowed understanding the modulation of bacterial recognition by the plants and revealed important aspects of plant immune responses^2^. Furthermore, additional investigations have confirmed that plant pathogenic bacteria exploit high flexibility in utilization of different kinds of sugar, nitrogen and phosphorus resources while adapting to the new environment *e*.*g*. bacterial plant pathogen *Pseudomonas syringae* pv. *tomato* specifically employs amino acid and sugar transporters to gain access to nutrients present in their environment. Subsequently, during infection processes of tomato plants, *P. syringae* pv. *tomato* is utilizing resources within the host, specifically directly from apoplast fluid^3^.

Pathogenic bacteria, to access the nutrients present in the plant tissues, colonize, invade and later on establish chronic infections within host plants. During the infection process, they enter plant tissues either through wounds or natural openings and occupy the apoplast of tissues or the xylem, where they multiply and spread. Specifically, phytopathogenic microorganisms cause damage and often impair plant growth and reproduction. On the other hand, to defend themselves against microbiological invasion, plants rely on two kinds of innate immunity, *i*.*e. via* pathogen triggered immunity (PTI)^4^ and effector-triggered immunity (ETI).

Hence, to achieve a compatible interaction, microorganisms at first need to overcome plant’s defenses that could abort the infection. Plant pathogenic bacteria, like *Pectobacterium parmentieri*, combine numerous strategies to accomplish that goal, *e*.*g*. they rely on quorum sensing system, hence the probability of weakening of plant defenses is higher due to high quantity of bacteria in plant environment^5^.

*P. parmentieri* is a pectinolytic, bacterium belonging to *Pectobacteriaceae* family (known as Soft Rot *Pectobacteriaceae*, SRP)^6^. It is a newly established species due to the transfer of all potato-originating isolates of *Pectobacterium wasabiae* to *P. parmentieri* sp. *nov*. (Ppa) on the basis of *in-silico* calculated DDH, gANI, and ANI values^7^. These Gram(-), rod-shaped bacteria are necrotrophs, that are able to destroy plant tissue components through the activity of PCWDE such as pectinases, cellulases, and proteases, further secreted via Type I or II secretion system^8,9^. Nonetheless, bacteria belonging to this species can exhibit a different level of activities of above-mentioned enzymes^10^. Pectinolytic bacteria, and among them *P. parmentieri*, require favorable environmental conditions to cause disease symptoms; however, they can reside inside plant tissues as endophytes for a long time^11^. Bacteria from the genus *Pectobacterium* are causative agents of soft rot in economically important crops such as potato, tomato or maize. Also, they are responsible for the blackleg disease, so far reported only on potato plants^11,12^. It is worth to mention that bacteria from the genus *Pectobacterium* have been included among 10 most important bacterial plant pathogens based on their economic impact^13^ since crop losses caused by phytopathogenic microorganisms can reach up to 20% of total yield^14^.

There is still limited knowledge regarding cascade of genes being expressed before and during infection process in *P. parmentieri*. It was previously reported, that initialization of infection progress is controlled by quorum sensing in closely related *Pectobacterium atrosepticum*^15^. Moreover, massive production of butanediol during plant infection by bacteria of the genera *Dickeya* and *Pectobacterium* was reported^16^. However, metabolic pathways important for promoting bacterial multiplication before and during plant infection connected with carbon and other compounds utilization in *P. parmentieri* have received little attention so far.

Given the complexity of bacterial-host relationships, they cannot be adequately investigated only by the means of classical microbiological and molecular methods; rather, the coupled use of Phenotype Microarrays™, computational, large-scale and systemic frameworks are advisable. Metabolic modeling is now a promising way to interpret puzzling, heterogeneous bacterial phenotypes^17^, especially those related to bacterial-host interaction^18^. Constraints-based approaches, and in particular Flux Balance Analysis (FBA)^19^ have been shown to infer growth phenotypes and are claimed to provide a systems biology view on multi-omics data, possibly allowing to predict physiological changes and evolution of bacterial populations^20,21^. Recently, genome-scale metabolic model (GEM) reconstruction and FBA have been used for deciphering the metabolic adaption of environmental microbes following ecological parameters variation^22^, ecological niche shift^23^, as well as for providing insights into the metabolic adaptation in human and bacterial plant pathogens^24,25^. To the best of our knowledge only in a few cases, FBA has been applied in understanding the metabolic adaptation of specific plant bacterial pathogens *e*.*g*. studies performed on *Rastonia solanacearum* showed that trade-off between virulence factor production and bacterial proliferation is controlled by the quorum-sensing-dependent regulatory protein *PhcA*^26^.

The aim of this work is the reconstruction of a highly curated metabolic model of the plant pathogenic bacterium *P. parmentieri* SCC3193 and the usage of this model to putatively identify the metabolic pathways relevant for *P. parmentieri* fitness in two different ecological niches, soil, and rhizosphere. We show that niche switching may lead to a metabolic reassessment in carbon-related pathways in *P. parmentieri* SCC3193 and we spot the core-set of essential genes in the two examined conditions. Moreover, we anticipate that the model itself will represent a valuable element which will pave the way to both, knowledge-base of strain’s biology and novel, applied technologies, like genetic engineering and synthetic biology experiments.

## RESULTS

### Reconstruction and validation of genome-scale metabolic model of *P. parmentieri* SCC3193

An initial draft model of *P. parmentieri* SCC3193 was obtained through the KBase server (http://kbase.us) then it has been reviewed and manually curated as described in the methods. The model was then augmented by mean of orthologous gene search in closely related *E. coli* strains. Following this approach, we hence expanded our reconstruction by adding 93 genes and 383 reactions from *E. coli* model^28^. Most of the added genes encode for transport reactions taking place in the membrane, where compounds are often modified and then transferred into the cytosol. This led to the necessity to add another compartment in *P. parmentieri* model, namely the periplasm, which improved descriptive model capability. After, model validation, and together further refinement, was carried-out by exploiting different approaches. Firstly, by comparing model’s performance to physiological data obtained in high throughput experiment with the use of PM Biolog plates. FBA was employed to test if the model could accurately predict the ability of *P. parmentieri* to produce biomass on utilized carbon sources. In particular, after verifying their presence in the metabolic reconstruction, we tested 91 compounds, previously used in PM experiment. The model was hence first gap-filled by iteratively correcting the inconsistencies between *in silico* predictions and PM outcomes. Where possible, missing reactions and transporters were added according to indications obtained from functional databases (MetaNetX^34^, Bigg^35^, Seed, KEGG^36^, and transportDB^37^) otherwise, gap-filling genes or reactions have been added and named after the missing components. Precisely, in twenty-two cases, the initial model’s prediction was refined thanks to experimental validations (further details on the predictions of metabolites usage, before and after this stage, are given in Supplementary Data SD1). At the end of this refinement round, the final *P. parmentieri* model was termed iLP1245 in accordance with the nomenclature standard^39^. It includes 1245 genes (covering ∼ 28% of the total number of coding sequences in the genome, 4449), 2182 reactions, and 2080 metabolites. A description of the model and the genetic features captured within are reported in Table 1.

**Table 1.**
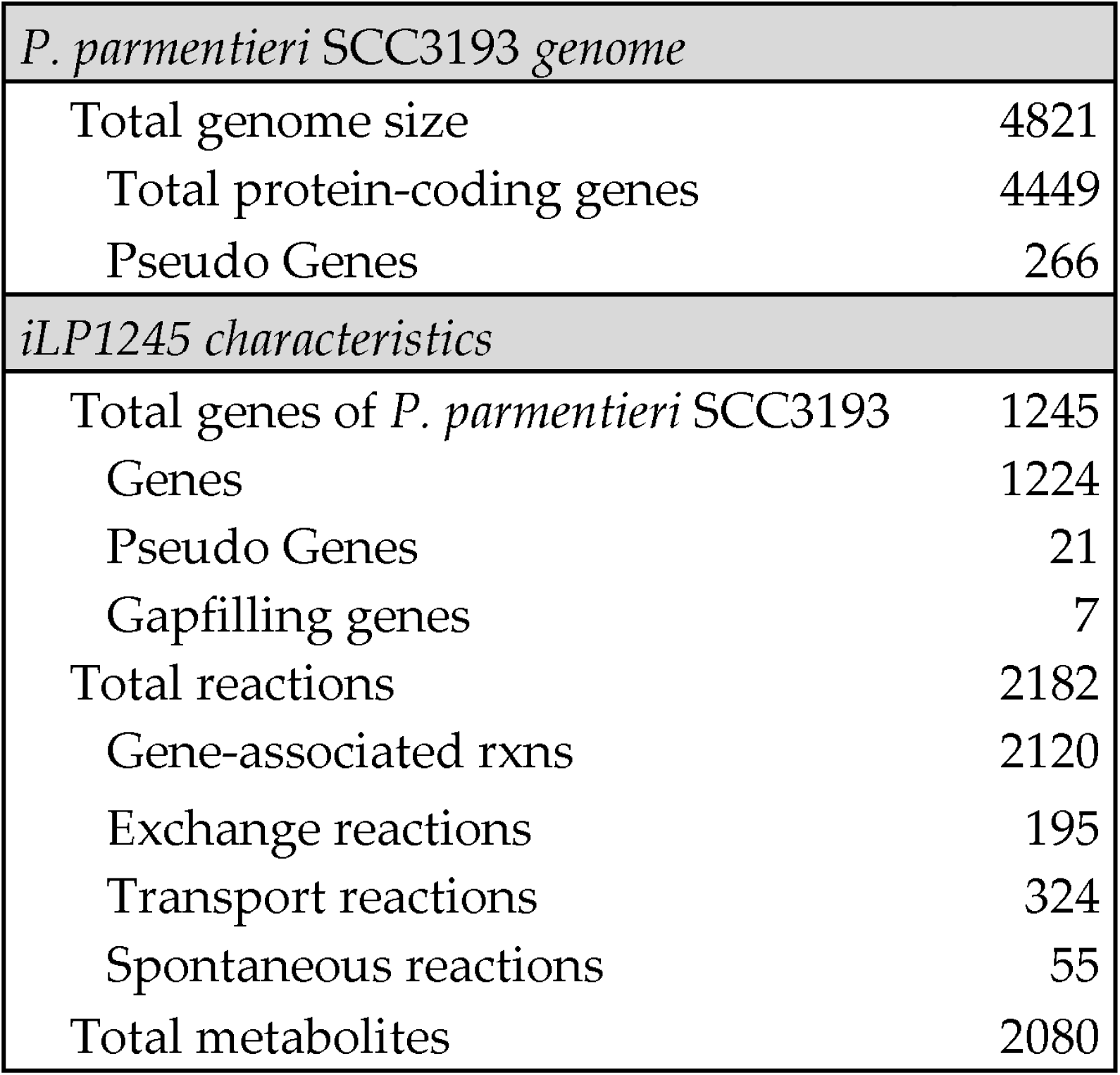
Properties of *P. parmentieri* SCC3193 genome and model

The refined model displayed agreement with PM in 83 out of 91 tested carbon substrates. As summarized in Figure 1, sensitivity, specificity, precision, accuracy, negative predictive value and F-score (calculated as described previously^38^, see materials and methods) reached very high scores, suggesting a good reliability of the model.

**Figure 1.**
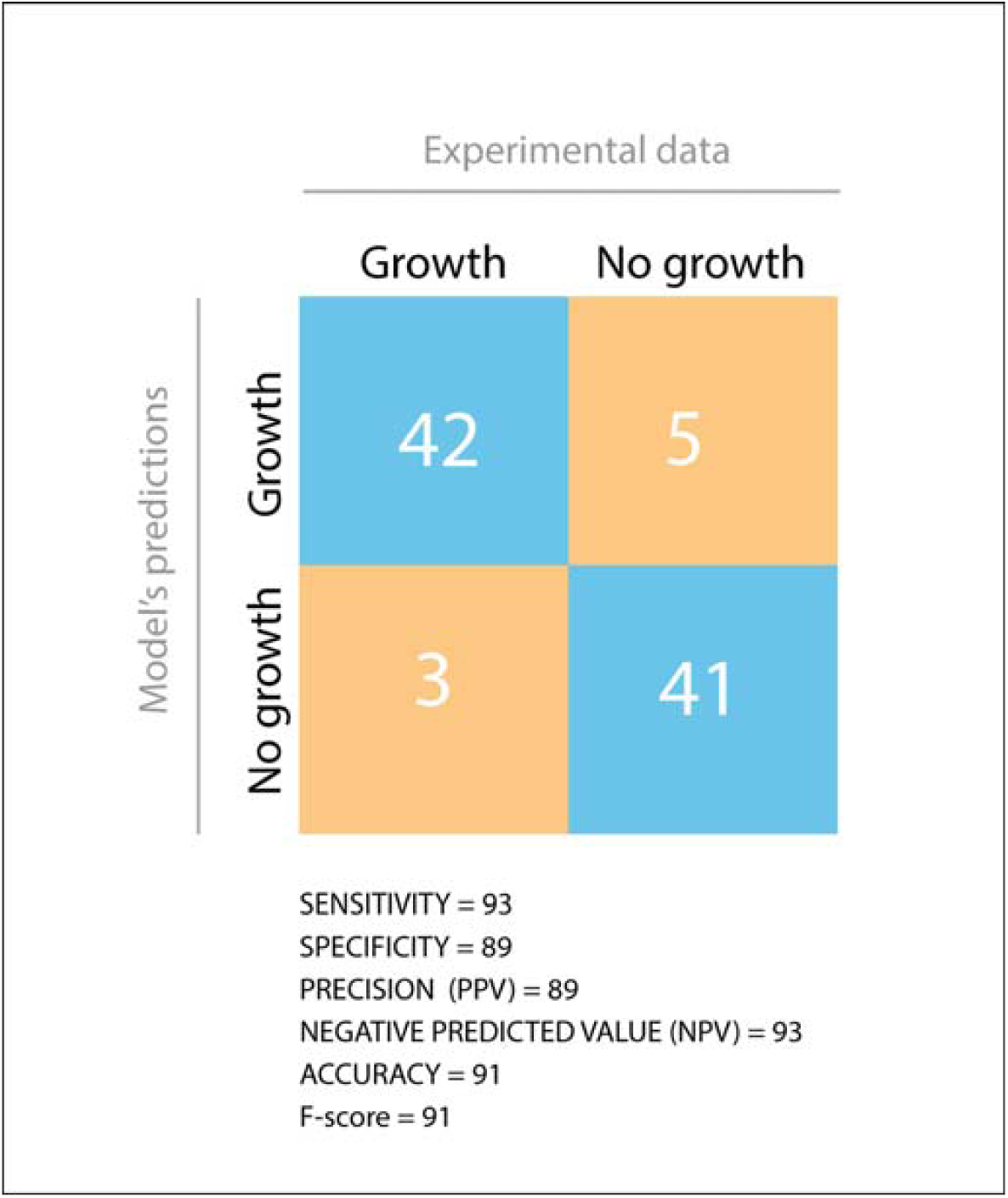
Comparison between Phenotype Microarray data and model’s predictions. TP = True Positive, TN = True Negative, FP = False Positive, FN = False Negative. Statistical parameters were calculated as described in materials and methods.

The Systems Biology Markup Language (SBML) file of the model was validated by the online SBML validator tool (http://sbml.org/Facilities/Validator/) and is available online as Supplementary Data SD2. All the metabolites embedded in the model were annotated by using identifiers.org and MIRIAM^40^ registry to facilitate model reuse and search strategies, by providing unique, unambiguous, perennial, standard-compliant and directly resolvable identifiers.

### Phenotypic characterization of *P. parmentieri* SCC3193 and metabolic model deep validation

Phenotypical profiling with the use of Biolog Plates PM1, PM2A, PM3 and PM4 revealed that *P. parmentieri* SCC3193 is able to utilize all common sugar components of plant cell walls at high levels *e*.*g*.: sucrose, tartaric acid, D-cellobiose, stachyose and most importantly pectin (Poly(1,4-alpha-D-galacturonide)) (SD3). Model’s predictions (SD5) are in agreement with *in vivo* obtained results. Precisely, this bacterium was very effectively exploiting D-Glucosamine and its derivatives (N-Acetyl-D-Glucosamine, N-Acetyl-D-Galactosamine) together with xanthine.

Cross-check of PM Microarray obtained results was achieved by applying EnVision™ analyses and measuring bacterial mass after 24 h of incubation in M9 media supplemented with randomly selected carbon sources: α-D-glucose, D-trehalose, and D-xylose.

Data obtained in the nutrients metabolic assay enabled to determine growth curves of *P. parmentieri* SCC3193 in M9 with three different carbon sources: α-D-glucose, D-trehalose, and D-xylose (Figure 2). Concerning bacterial growth in α-D-glucose, we observe rapid logarithmic phase and plateau of bacterial culture (Figure 2). Interestingly, in case of D-trehalose and D-xylose a longer lag phase occurs followed by an intermediate logarithmic phase. Comparison of EnVision™ obtained results to models’ predictions revealed agreement in case of all 3 tested carbon sources (Figure 2).

**Figure 2.**
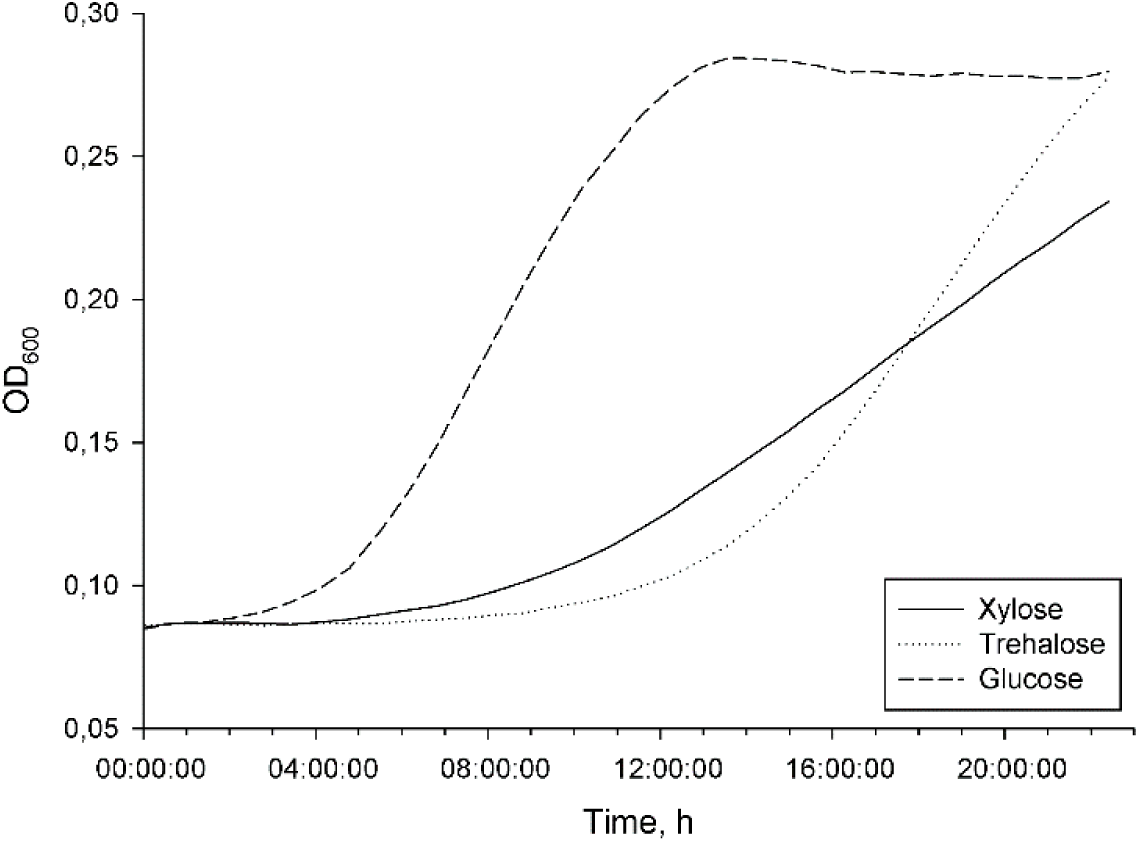
Growth curves of *P. parmentieri* SCC3193 in M9 supplemented with different carbon sources.

### Differential metabolic adaptation to soil and rhizosphere environment

By analyzing flux changes in response to simulated environmental conditions (*e*.*g*. M9, soil, rhizosphere) it is possible to observe whether any significant metabolic rewiring occurs. Loopless FBA and loopless FVA (see materials and methods) were used in order to estimate niche-specific metabolic adaptations in soil and rhizosphere. The results obtained through FBA were cross-checked with FVA (see Supplementary Data SD3 for results). Possible fluxes variation occurring during the transition from soil to rhizosphere was interpreted as a metabolic shift between the two niches. The magnitude of the variation was described according to different cutoff, *i*.*e*. as a variation greater than 10%, 20%, 30% 40% and 50% of the initial flux value. Results indicate that the number of the reactions which fluxes changed significantly is slightly affected by the stringency of the cutoff, vouching for robustness of the analysis. Accordingly to the previous report^23^, we focused only on the results for which variation is higher than 50% in rhizosphere compared to soil. Finally, the total amount of reactions taken into account was 208, corresponding to ∼ 10% of those embedded in the reconstruction. These were further classified as reactions whose flux increase or decrease and reactions turned on or off during the environmental stress (see Figure 3). Based on that classification, it appears that several pathways keep being active in both examined conditions, though recruiting a different set of reactions. The main shift seems to occur in sugars metabolic pathways, tracking a switch from pentose phosphate and hexose to amino sugars metabolism. Also, some peculiar systems seem to be turned on in concomitance of such niche change, as nitrogen, butanoate, galactose, and propanoate metabolism, and biosynthetic pathways, including those of steroids and folate, attesting a specific adaptive response of *P. parmentieri* SCC3193 to this environmental (nutritional) switch.

**Figure 3.**
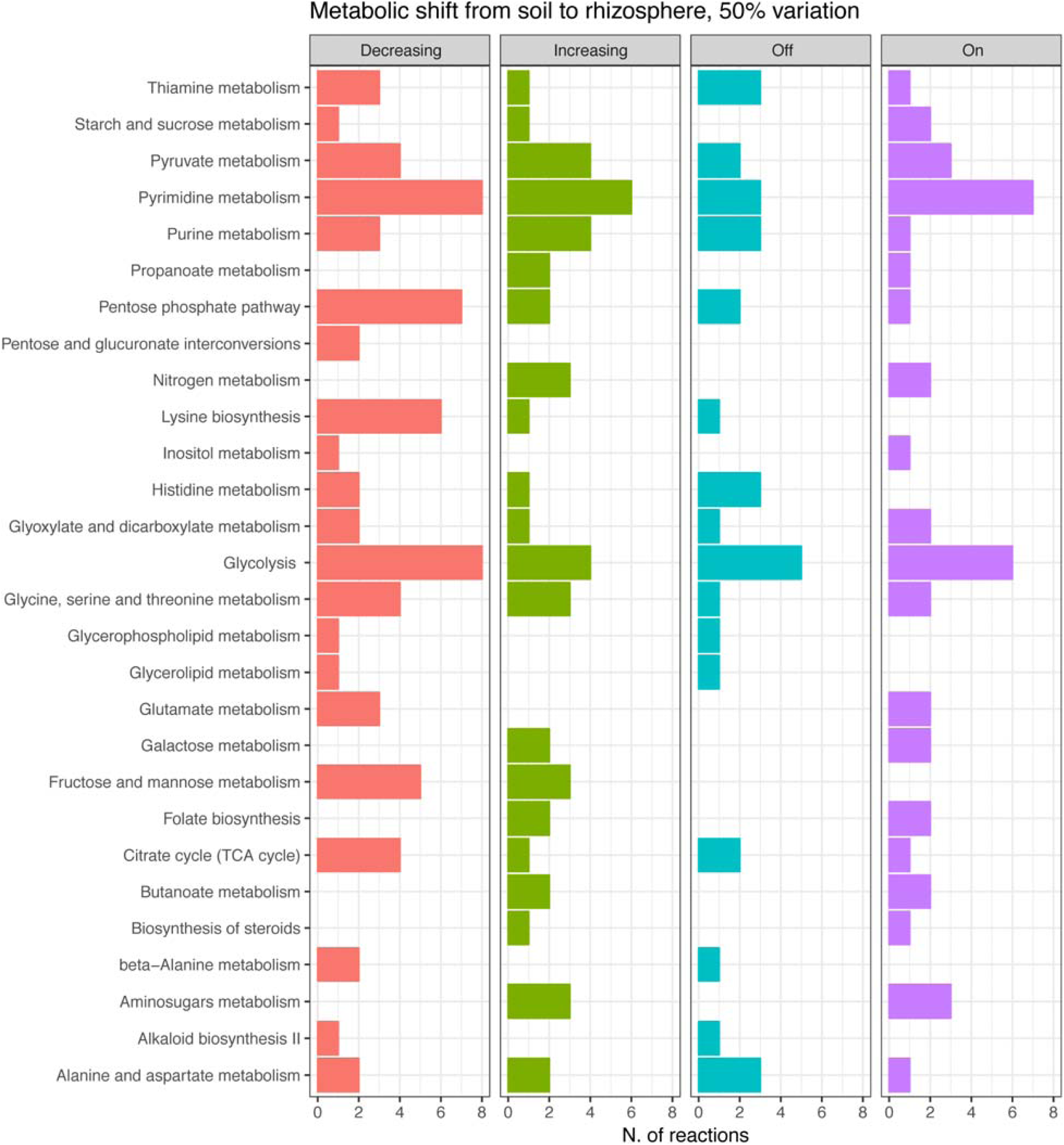
Biochemical reactions which fluxes change, while bacterial transition from soil to the rhizosphere.

### *In silico* genes deletion provides insight into fitness relevance of metabolic modules

Simulated single genes deletion is a very powerful *in silico* method, as it not only enables to estimate gene’s knock-out fatality but also, allows predictions on a genome-scale. We performed such simulations on iLP1245 with two different methods (see materials and methods), FBA and MOMA. As shown in Figure 4, both methods labeled the same number of genes as essential, according to the three media examined, hence from now on we will refer only to FBA outcomes. Specifically, the number of essential genes found is 251 in M9 medium, 250 in soil and 245 in the rhizosphere. The locus tag of such genes, alongside with their corresponding encoded protein, are reported in Supplementary Data SD4. The Venn’s diagram in Figure 5 shows the overlap among the three different conditions. Results indicate that a huge core of genes is likely to be mandatory in all tested *in silico* conditions, while only small number stand-out as essential just in one or two out of the three tested conditions. Indeed, in *in silico* gene deletions predicted the gene W5S_RS13875 encoding for DNA starvation/stationary phase protection protein (WP_014700482.1) as specifically required during growth in soil and rhizosphere, the gene W5S_RS15765 encoding for an ammonium transporter (WP_014700821.1) essential in M9 and rhizosphere, while eight genes appear to be specifically essential during growth in soil and M9 but not in rhizosphere. Seven of them belong to the thiamine and sulfur metabolism pathway: W5S_RS00965 cystathionine gamma-synthase, WP_014698476.1), W5S_RS01140 (thiazole synthase ThiG, WP_012822036.1), W5S_RS01145 (sulfur carrier protein ThiS, WP_014698484.1), W5S_RS01150 (adenylyltransferase ThiF, WP_014698485.1), W5S_RS01155 (thiamine phosphate synthase ThiE, WP_014698486.1), W5S_RS01160 (phosphomethylpyrimidine synthase ThiC, WP_014698487.1), W5S_RS05940 (hydroxymethylpyrimidine / phosphomethylpyrimidine kinase ThiD, WP_014698998.1), while the last one, W5S_RS18250 (diaminopimelate decarboxylase, WP_043899153.1) is an enzyme involved in secondary metabolite production.

**Figure 4.**
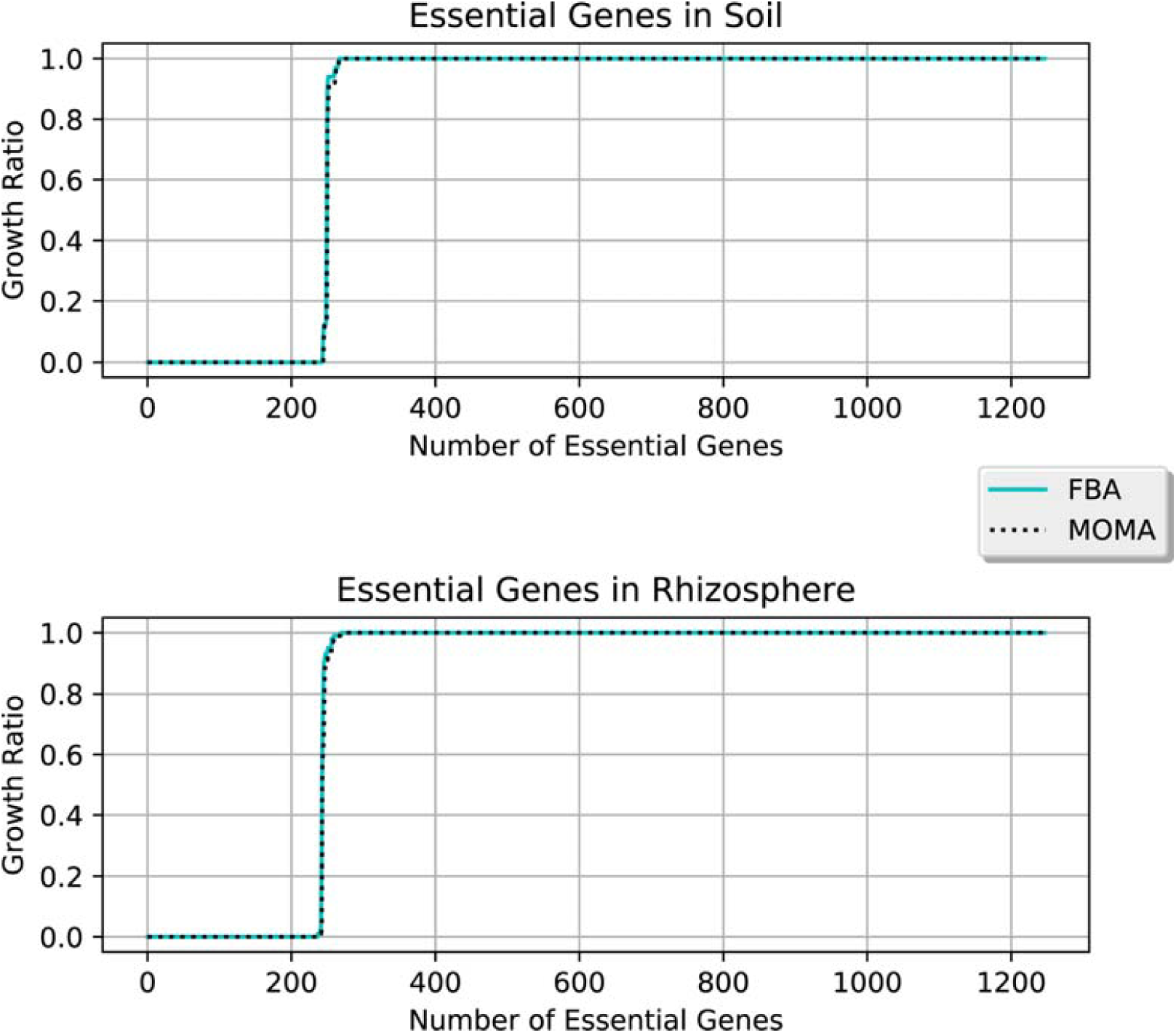
GRratio value for each gene deletion in rhizosphere and soil media according to FBA and MOMA predictions.

**Figure 5.**
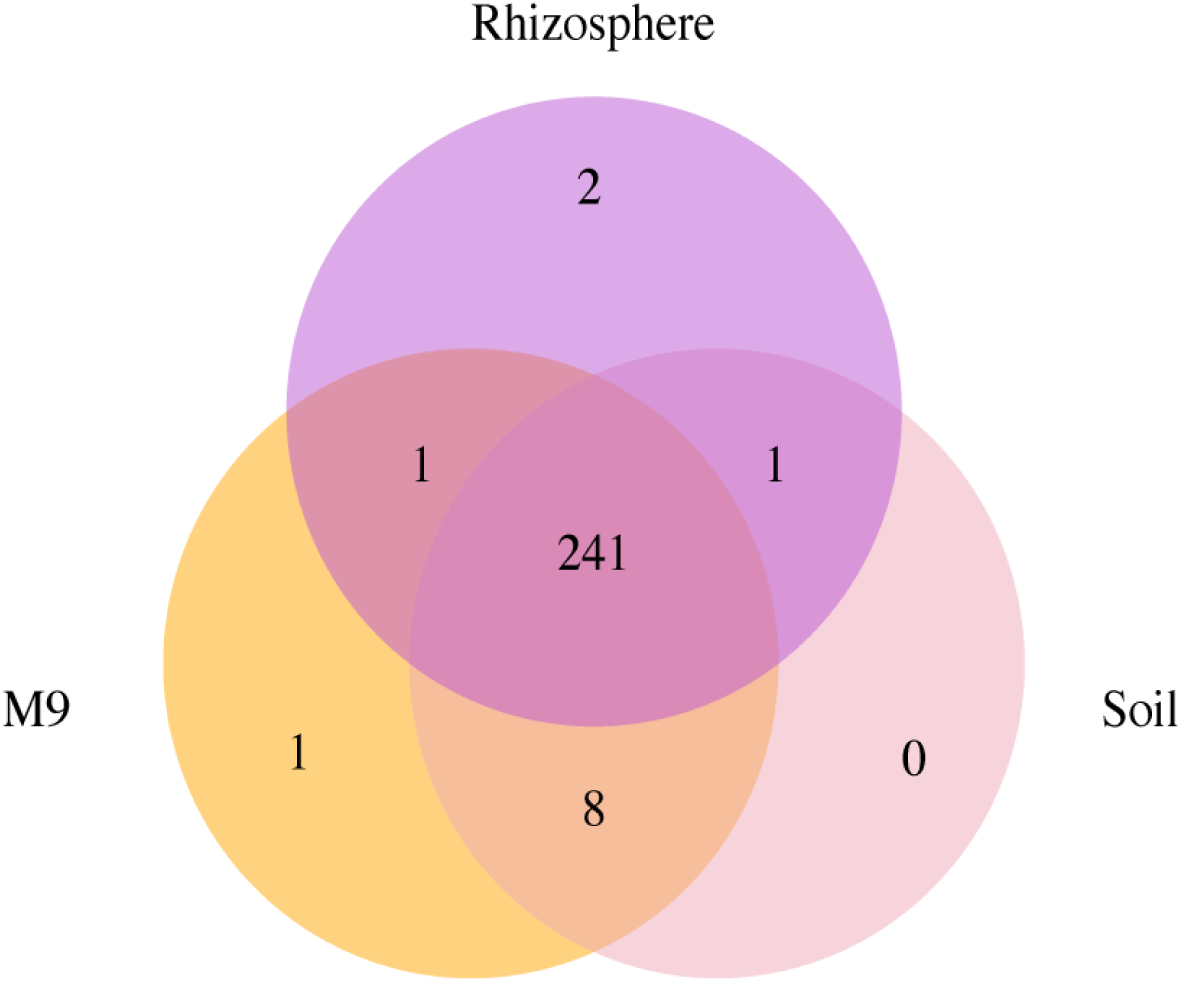
Venn’s diagram showing the amount of shared/unique essential genes for each of the examined conditions (M9, rhizosphere, and soil).

Interestingly, the simulated rhizosphere growth medium contains thiamine which, conversely, is absent in soil and M9.

Moreover, three condition-specific genes have been predicted, one during growth in M9 medium, W5S_RS19605 encoding for class II fructose-bisphosphate aldolase (WP_014701539.1) and two during growth in the rhizosphere, W5S_RS19610 and W5S_RS06525, respectively encoding for a phosphoglycerate kinase (WP_005973111.1) and a long-chain fatty acid transporter (WP_014699104.1).

## DISCUSSION

Plant-microbe interactions have been under intensive investigation in recent years, regardless of the nature of this communication: pathogenic or symbiotic. Severity to understand this interplay is connected with the complexity of the environment in which bacteria persist: soil, rhizosphere or plant tissues. Metabolic modeling allows predicting and examining biochemical reactions involved in adaptation to above-mentioned ecological niches as well as predicting the phenotypic outcomes of gene deletions. In this paper, for the first time we report a high throughput experimental validation on metabolic capability of *P. parmentieri* SCC3193, and also a highly curated metabolic model of this plant pathogenic bacterium (iLPI245).

iLPI245 is highly reliable, as *in silico* obtained results overlap in 91% with experimentally obtained data on carbon utilization phenotypes, a value that perfectly fits with the currently accepted standard for genome-scale metabolic reconstructions^41,42^. For example, a previously described genome-scale metabolic reconstruction of *Pectobacterium aroidearum* PC1, a monocotyledonous plant pathogenic bacterium, showed the agreement of 80.4% between *in silico* simulations and Phenotype Microarray™ (Biolog) experiments^25^. This difference in accuracy of the models was probably related to the fact, that the metabolic model of *P. aroidearum* PC1 was constructed on the template of the latest version of *E. coli* K12 MG1655 metabolic model iJO1366, whereas in our case *E. coli* genome was used only for biomass estimation, and identification of orthologues genes, in consequence producing a more *P. parmentieri*-specific model.

The iLP1245 model of *P. parmentieri* SCC3193 was then used to assess the relative importance of single metabolic pathways and genes in adaptation to growth under laboratory (M9 medium) and *in silico* simulated field conditions (soil and rhizosphere). Soil and rhizosphere conditions were chosen since they represent two key environments were *P. parmentieri* is abundant, and for which the information on chemical composition is readily available and reliable^23^. While performing FBA on *P. parmentieri* SCC3193, we observed a shift in sugars metabolic pathways while conditions were changed from soil to rhizosphere. Namely, switch from pentose phosphate and hexose to amino sugars metabolism, which are supposed to be more abundant in rhizosphere^43^. We can consequently suggest, that adapting to rhizosphere environment involves utilization of this latter carbon compounds. Interestingly, a down-regulation of genes expression involved in amino sugars and nucleotide sugar metabolism has been associated with starvation stress response in *P. atrosepticum*^44^. Then, our prediction of metabolic fluxes implies, that rhizosphere represents a rich environment for *P. parmentieri*, where these bacteria can thrive. Moreover, butanoate, propanoate, steroid metabolism and, folate biosynthetic pathways are turned on while we *in silico* shift environment from soil to the rhizosphere. These compounds are precursors for volatiles compounds (VOCs) production or are comprised in VOC metabolic pathways; *e. g*. butanoate is an ester of butyric acid, which is among the most frequently secreted compounds^16,45^. We can hypothesize that interbacterial communication and possibly plant-bacteria interaction could be mediated by VOCs, which may then have a role in pathogenesis and later on, in developing strategies for biocontrol. All those volatiles are strictly connected either with encountering other bacteria growing in rhizosphere or with virulence of plant pathogenic bacteria^16,45^.

These data were compared with those obtained from the plant symbiotic nitrogen-fixing bacterium *Sinorhizobium meliloti*^23^. We observed that only 10% of reactions changed flux (at least by 50%) in *P. parmentieri*, while the shift from soil to rhizosphere conditions changed the flux of more than 20% of reactions (including reactions which reversed direction) in *S. meliloti*, though the number of reactions present in the two models (iLP1245 and iGD1575) was similar (2182 and 1.825 reactions, respectively). Moreover, in *S. meliloti* ∼ 13% of active reactions were specific to just one of the environments, while only the 5.3% of active reactions were environment specific in *P. parmentieri*. We can hypothesize that the smaller and more compact genome of *P. parmentieri* compared to *S. meliloti* (4449 *vs*. 6204 protein-coding genes for *P. parmentieri* and *S. meliloti*, respectively, including a multipartite genome organization in the latter species) allows a reduced metabolic redundancy in *P. parmentieri* compared to *S. meliloti*^46^ and a more generalist *vs*. specialist metabolic network (*i*.*e*. most reactions are not changing while environment fluctuates). To test this hypothesis, we performed MOMA and FBA simulations of gene deletions (see Supplementary Data SD4). Both analyses revealed that *P. parmentieri* SCC3193 possesses an essential gene core composed of 241 metabolic genes, for growth in both soil and the rhizosphere. Here, a set of 8 genes only was found to be essential in soil but not in rhizosphere. This is in large contrast to *S. meliloti*, where 66 genes were found as essential for growth in the same simulated rhizosphere environment^23^, and supports the previously proposed hypothesis of a robust and versatile metabolic network of *P. parmentieri* SCC3193, which may allow the strain to rapidly accommodate with relevant changes in environmental nutrient sources. Additionally, we can assume that compact core of essential genes important for *P. parmentieri* is a reason why bacteria from this species are able to persist on plant residuals without interacting with host plant (potato) for long periods of time, and as well are cosmopolites in the environment^12^.

To summarize our findings, we established a functional metabolic model of plant pathogenic bacterium *P. parmentieri* SCC3193, showing an inherent robustness of the metabolic network, which can easily accommodate for nutrient variability when moving from soil to rhizosphere growth conditions. Such robustness may imply that several, still unknown, plant species (together with their soil and rhizosphere) can be a reservoir for this pathogenic bacterium.

## MATERIALS AND METHODS

### Metabolic network reconstruction and model refinement

A draft metabolic model was build using the KBase Narrative Interface (www.kbase.com), and later it was expanded based on the comparison of identical functions (orthologs genes) in the closely related *Escherichia coli* K-12 MG1655 (through a Bidirectional Best Hit approach, inParanoid^27^), for which a reliable metabolic reconstruction is available^28^.

Further model refinement was then performed through experimental validation, *i*.*e*. by iteratively comparing model’s growth outcomes to the real strain’s growth during Phenotype Microarrays experiment (Omnilog™). Additional information on the comparison is reported in Supplementary Data SD1.

### Metabolic modeling

Metabolic modeling was performed using COBRApy Toolbox version 0.6.1^29^ and the Gurobi 7.0.2 solver (www.gurobi.com). Scripts enabling to run all the *in silico* analysis performed in this work are available in Supplementary Information SI1. No comprehensive description of the macromolecular composition of the *P. parmentieri* biomass is available in the literature. However, such data are available for *E. coli*^28^, therefore we approximated the *P. parmentieri* gross biomass composition to that one of this closely related species. The complete biomass composition is given in Supplementary Information SI2. The biomass reaction was set as the objective function for growth in all the experiments performed in this work.

### Bacterial strains and culture conditions

Bacterial strain used in this study is *P. parmentieri* reference strain SCC3193 isolated from potato tuber in Finland^7,30^. For high-throughput phenotypic characterization bacteria were grown on TSA medium at 28°C for 24 h. For EnVision™ experiment bacteria were first grown in LB at 28°C for 24 h with constant agitation (120 RPM), later on in M9 for 24 h with constant agitation (130 RPM).

### Experimental high-throughput phenotypic characterization on *P. parmentieri* SCC3193

For high-throughput phenotypic characterization of *P. parmentieri* SCC3193 Biolog Plates PM1, PM2A, PM3 and PM4 were utilized. Overnight bacterial culture was transferred from TSA medium to 5 ml of 0.85% NaCl and bacterial suspension was adjusted to OD_600_ equaling 0.1 Later, 1 ml of bacterial suspension was transferred to 11 ml of Minimal Salts medium (M9-C: % Na_2_HPO_4_, 0.3% KH_2_PO_4_, 0.05% NaCl, NH_4_Cl, 0.005% Yeast Extract) supplemented with 120 µl of Biolog A dye. To inoculate wells in PM plates 100 µl of described bacterial suspension was used. The measurement was carried out in OmniLog™ for 46h. Results were analyzed with DuctApe^31^. All the results are reported in Supplementary Data SD5.

Nutrients metabolic assay with the use of EnVision™ plate reader was performed to cross-check high-throughput phenotypic characterization. M9 media supplemented with 20% of selected carbon sources: α-D-glucose, D-xylose, and D-trehalose were prepared. *P. parmentieri* SCC3193 was grown in LB medium overnight at 28°C with constant agitation (120 RPM). Afterwards, overnight bacterial cultures were centrifuged and washed twice in sterile Ringer Buffer and later OD_600_ of inoculum was adjusted to 0.1. To establish growth curves 50 µl of inoculum was transferred to 450 µl of M9 supplemented with different carbon sources. Bacteria were cultured with agitation set at 120 RPM in 28°C in 24-well plate in EnVision™ plate reader. Optical density measurement at 600 nm was performed every 30 min.

### OmniLog™ data processing and analysis

PM1 and PM2A obtained data analysis was performed with DuctApe^31^. Activity index (AV) values were calculated following subtraction of the value obtained for blank well from that of inoculated wells, whereas plots of the growth curves are of the unblanked data. Bacterial growth with each compound was considered positive if the AV value was ≥ 3. Growth phenotypes were defined as negative if the AV value was ≤ 2, and following a manual inspection of the unblanked curves. All the results are reported in Supplementary Data SD6.

### *In silico* environmental representations

*In silico* representations of the nutritional composition of the soil, and rhizosphere were derived from a previously published paper^23^. The composition of M9, soil, and rhizosphere media is reported in SI2.

### Flux Balance Analysis and Flux Variability Analysis

Flux distribution predictions were assessed by performing FBA in M9, soil and rhizosphere media. Moreover, to avoid *in silico* artifacts, such as loops (which looks unreal to happen *in vivo*), we performed loopless-FBA, which instead identifies the closest (to the reference) thermodynamically consistent flux state avoiding loops.

Moreover, since FBA only predicts one flux distribution among all possible solutions, we also performed loopless-FVA (Flux Variability Analysis) and filtered out those reactions whose fluxes were outside the following criteria:

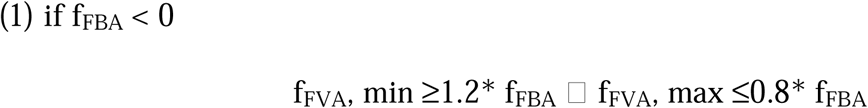

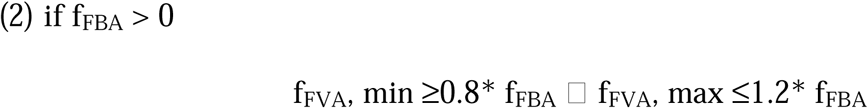

Loopless FBA and FVA predictions for each reaction can be retrieved in Supplementary data SD3 or by running Supplementary information SI1.

### Model’s predictive value estimation

M9 growth medium was simulated *in silico, i*.*e*. by constraining the lower bound of import reactions for each of the compounds present in the medium (as reported in SI2). When reproducing PM experiment *in silico*, model’s performances were considered as true positives (TP) if growth was obtained both *in silico* and *in vivo*, true negatives (TN) in case of non-growth both *in silico* and *in vivo*, false positives (FP) if growth was obtained *in silico* but not *in vivo* and false negatives (FN) if *vice versa*.

Reliability of the obtained predictions was then estimated according to the following parameters:

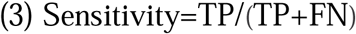

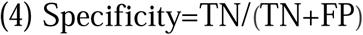

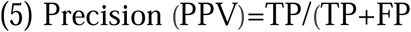

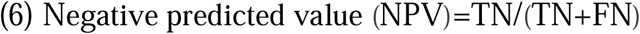

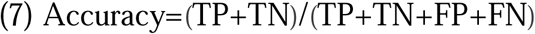

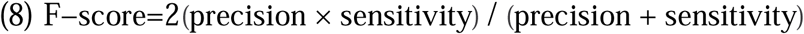

### Single gene deletion analysis

Using genome-scale metabolic networks (GEMs) genes knockout can be simulated to identify those genes whose removal is likely to impair the organism’s growth. Specifically, it is possible to simulate mutants by deleting each gene included in the metabolic reconstruction and testing the predicted effects on the microbe’s growth. Through this strategy, it is possible to calculate the growth ratio (GR_ratio_) between the growth rate of the mutant model (*μ*KO) and the one of the wild-type (*μ*WT) as:

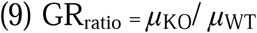

This measure can provide hints on the essentiality of the knocked out gene. In particular, in our work the knocked out gene was considered essential when GR_ratio_ < 0.9. Furthermore, when 0.9 < GR_ratio_ < 1 and GR_ratio_ = 1 we considered the knocked out gene as semi-essential and essential, respectively. Minimization of Metabolic Adjustment (MOMA^32^) and FBA were the algorithms chosen to perform such analyses.

### COG analyses

The WebMGA webserver^33^ was used to provide functional Cluster of Orthologous Genes (COG) annotations (p-value cutoff of 0.001) to each gene in the model. The COG annotation for each gene associated with variable reactions during the transition between two niches was extracted from the WebMGA server. Biases were determined after standardizing by the number of genes in each class of variable genes. Statistical significance was determined using Pearson’s Chi-squared tests. The complete list of COG annotations is available as Supplementary Data SD7.

## ACKNOWLEDGEMENTS

This work was supported by the National Science Centre in Poland via grant no. 2014/14/M/NZ8/00501 awarded to EL and by the University of Gdansk in Poland via grant no. 538-M031-B187-16 awarded to SZ.

## AUTHOR CONTRIBUTIONS STATEMENT

E.L and A.M. conceived the work, L.P. reconstructed the model and performed the computational experiments, S.Z. and F.D. conducted the wet-lab experiments, M.F., A.M., L.P.,L.G. and S.Z. analyzed the results. L.P. and S.Z. draft the manuscript. All authors reviewed the manuscript.

## SUPPLEMENTARY INFORMATION

The authors declare that they have no competing interests.

## SUPPLEMENTARY FILES

**Supplementary Information 1**. Script enabling to perform all the analysis in the manuscript

**Supplementary Information 2**. Includes the Supplementary Text, Supplementary Tables, and Supplementary Figures. File type: PDF document

**Supplementary Data 1**. Prediction on usage of metabolites before and after gap filling process. File type: Excel document

**Supplementary Data 2**. The sbml file of the model. File type: XML formatted file

**Supplementary Data 3**. Reactions Flux during growth in soil *versus* the rhizosphere according to loopless FBA and loopless FVA. File type: Excel document

**Supplementary Data 4**. All essential genes found with their encoded protein and function. Growth ratio calculated with FBA and MOMA is shown.

**Supplementary Data 5**. All Phenotype MicroArray™ data generated in this study and plots representing results from Phenotype MicroArray™ analysis with the use of DuctApe program. File type: Compressed zip archive

**Supplementary Data 6**. The raw Phenotype MicroArray™ data, in the form of .csv files, obtained in this study. File type: Compressed zip archive

**Supplementary Data 7**. The COG annotations for all genes included in the model, as generated by WebMGA. File type: Excel document

