## Supplementary Materials for "Metabolic modeling of *Pectobacterium parmentieri* SCC3193 provides insights into metabolic pathways of plant pathogenic bacteria"

Supplementary Informations SI2.

**Supplementary Table 1**. M9, soil, and rhizosphere media composition.

| Compound Name | Exchange Reaction | LB in soil | LB in rhizosphere | LBin M9 |
|---|---|---|---|---|
| $H_2O$ | EX_cpd00001_e0 | -15 | -15 | -10 |
| $O_2$ | EX_cpd00007_e0 | -15 | -15 | -10 |
| Phosphate | EX_cpd00009_e0 | -15 | -15 | -10 |
| $CO_2$ | EX_cpd00011_e0 | -15 | -15 | 0 |
| Ammonia | EX_cpd00013_e0 | -7.5 | -7.5 | -10 |
| L-glutamate | EX_cpd00023_e0 | 0 | -0.0283302 | 0 |
| D-glucose | EX_cpd00027_e0 | -0.61972444 | -0.04098397 | 0 |
| $Mn_2$ | EX_cpd00030_e0 | -15 | -15 | -10 |
| Glycine | EX_cpd00033_e0 | -0.0068175 | -0.00693094 | 0 |
| $Zn_2$ | EX_cpd00034_e0 | -15 | -15 | -10 |
| L-alanine | EX_cpd00035_e0 | -0.02780553 | -0.00823049 | 0 |
| Succinate | EX_cpd00036_e0 | -0.0056245 | -0.12240603 | 0 |
| L-lysine | EX_cpd00039_e0 | 0 | -10 | 0 |
| L-aspartate | EX_cpd00041_e0 | 0 | -0.03205557 | 0 |
| Sulfate | EX_cpd00048_e0 | -15 | -15 | -10 |
| L-arginine | EX_cpd00051_e0 | -0.0068175 | -0.00948672 | 0 |
| L-serine | EX_cpd00054_e0 | 0 | -0.01004986 | 0 |
| $Cu^{2+}$ | EX_cpd00058_e0 | -15 | -15 | -10 |
| $Ca^{2+}$ | EX_cpd00063_e0 | -15 | -100 | -10 |
| L-ornithine | EX_cpd00064_e0 | -0.0068175 | -0.00831712 | 0 |
| $H^+$ | EX_cpd00067_e0 | -15 | -15 | -10 |
| L-tyrosine | EX_cpd00069_e0 | -0.0068175 | -0.00233919 | 0 |
| Sucrose | EX_cpd00076_e0 | 0 | -0.02049199 | 0 |
| L-cysteine | EX_cpd00084_e0 | -0.0068175 | 0 | 0 |
| $Cl^-$ | EX_cpd00099_e0 | -15 | -15 | -10 |
| Glycerol | EX_cpd00100_e0 | 0 | 0 | -10 |
| Biotin | EX_cpd00104_e0 | -15 | -15 | 0 |
| D-ribose | EX_cpd00105_e0 | -0.01862144 | 0 | 0 |
| L-leucine | EX_cpd00107_e0 | -0.03596182 | -0.00303228 | 0 |
| D-galactose | EX_cpd00108_e0 | -0.25290619 | -0.18317325 | 0 |
| L-histidine | EX_cpd00119_e0 | -0.0068175 | -0.00506825 | 0 |
| L-proline | EX_cpd00129_e0 | -0.01102953 | 0 | 0 |
| L-malate | EX_cpd00130_e0 | -0.03649016 | -0.79413596 | 0 |
| D-mannose | EX_cpd00138_e0 | -0.2540567 | -0.05436649 | 0 |

| | | | | |
|---|---|---|---|---|
| $CO_2$ | EX_cpd00149_e0 | -15 | -15 | -10 |
| Xylose | EX_cpd00154_e0 | -0.21280795 | -0.04600242 | 0 |
| L-valine | EX_cpd00156_e0 | -0.02928849 | -0.00337883 | 0 |
| L-threonine | EX_cpd00161_e0 | -0.0068175 | -0.00961667 | 0 |
| $K^+$ | EX_cpd00205_e0 | -15 | -15 | -10 |
| Nitrate | EX_cpd00209_e0 | -7.5 | -7.5 | 0 |
| L-arabinose | EX_cpd00224_e0 | -0.28889705 | -0.26974143 | 0 |
| $Mg^{2+}$ | EX_cpd00254_e0 | -15 | -15 | -10 |
| spermidine | EX_cpd00264_e0 | 0 | -10 | 0 |
| Thiamine | EX_cpd00305_e0 | 0 | -10 | 0 |
| L-isoleucine | EX_cpd00322_e0 | -0.0199273 | -0.00186269 | 0 |
| D-raffinose | EX_cpd00382_e0 | | -0.01024599 | 0 |
| L-rhamnose | EX_cpd00396_e0 | -0.13913636 | -0.02927426 | 0 |
| Trans-4-hydroxy-l-proline | EX_cpd00851_e0 | 0 | -0.26974144 | 0 |
| $Na^+$ | EX_cpd00971_e0 | -15 | -15 | -10 |
| octadecanoate | EX_cpd01080_e0 | 0 | -10 | 0 |
| Stachyose | EX_cpd01133_e0 | 0 | -0.03073798 | 0 |
| tetradecanoate | EX_cpd03847_e0 | 0 | -10 | 0 |
| $Fe^{2+}$ | EX_cpd10515_e0 | -15 | -15 | -10 |
| $Fe^{3+}$ | EX_cpd10516_e0 | -15 | -15 | -10 |

**Supplementary Table 2.** List of genes-protein-function embedded in the model.

| LOCUS TAG | PROTEIN | FUNCTION |
|---|---|---|
| W5S_RS11060 | WP_043899042.1 | L-arabinose isomerase |
| W5S_RS03460 | WP_012822536.1 | dihydropteroate synthase |
| W5S_RS19660 | WP_014701549.1 | glutathione synthase |
| W5S_RS05190 | WP_033071956.1 | hypoxanthine phosphoribosyltransferase |
| W5S_RS03645 | WP_014698749.1 | lactaldehyde reductase |
| W5S_RS10745 | WP_014699889.1 | nicotinamidase/pyrazinamidase |
| W5S_RS14305 | WP_014700565.1 | biotin synthase |
| W5S_RS14775 | WP_014700661.1 | MFS transporter |
| W5S_RS12660 | WP_014700262.1 | beta-galactosidase |
| W5S_RS08210 | WP_014699430.1 | Evolved beta-D-galactosidase subunit alpha |
| W5S_RS06020 | WP_014699012.1 | beta-galactosidase |
| W5S_RS03025 | WP_012822358.1 | serine acetyltransferase |
| W5S_RS21915 | WP_014701938.1 | serine O-acetyltransferase |
| W5S_RS09490 | WP_014699662.1 | cytidylate kinase |
| W5S_RS14685 | WP_014700645.1 | anaerobic C4-dicarboxylate transporter DcuC |
| W5S_RS01995 | WP_014698568.1 | TRAP transporter substrate-binding protein DctP |

| | | |
|---|---|---|
| W5S_RS18995 | WP_014701427.1 | undecaprenyldiphospho-muramoylpentapeptide beta-N- acetylglucosaminyltransferase |
| W5S_RS18800 | WP_014701388.1 | isochorismatase |
| W5S_RS13670 | WP_014700443.1 | isochorismatase |
| W5S_RS19065 | WP_014701438.1 | long-chain fatty acid--CoA ligase |
| W5S_RS10580 | WP_014699859.1 | long-chain-fatty-acid--CoA ligase |
| W5S_RS04990 | WP_014698887.1 | glutamate--cysteine ligase |
| W5S_RS12570 | WP_014700244.1 | enoyl-[acyl-carrier-protein reductase |
| W5S_RS06510 | WP_014699101.1 | malate:quinone oxidoreductase |
| W5S_RS05735 | WP_014698955.1 | nucleoside-diphosphate kinase |
| W5S_RS20535 | WP_014701684.1 | phosphoenolpyruvate carboxykinase (ATP) |
| W5S_RS18875 | WP_014701403.1 | pyruvate dehydrogenase (acetyl-transferring), homodimeric type |
| W5S_RS21180 | WP_014701804.1 | undecaprenyl-phosphate alpha-N-acetylglucosaminyl 1-phosphate transferase |
| W5S_RS14675 | WP_014700643.1 | undecaprenyl-phosphate alpha-N-acetylglucosaminyl 1-phosphate transferase |
| W5S_RS12615 | WP_014700253.1 | alpha-glucosidase |
| W5S_RS12765 | WP_014700280.1 | aconitate hydratase |
| W5S_RS18805 | WP_014701389.1 | aconitate hydratase B |
| W5S_RS14575 | WP_014700619.1 | dTDP-4-dehydrorhamnose 3,5-epimerase |
| W5S_RS21365 | WP_014701831.1 | multifunctional fatty acid oxidation complex subunit alpha |
| W5S_RS06540 | WP_014699107.1 | multifunctional fatty acid oxidation complex subunit alpha |
| W5S_RS15970 | WP_014700851.1 | tRNA guanosine(34) transglycosylase Tgt |
| W5S_RS12955 | WP_014700316.1 | catalase/peroxidase HPI |
| W5S_RS13540 | WP_014700418.1 | malonyl CoA-acyl carrier protein transacylase |
| W5S_RS20855 | WP_005969274.1 | aspartate-semialdehyde dehydrogenase |
| W5S_RS17950 | WP_043899149.1 | DNA ligase (NAD(+)) LigA |
| W5S_RS00005 | WP_005976670.1 | chromosomal replication initiation protein DnaA |
| W5S_RS00010 | WP_005976672.1 | DNA polymerase III subunit beta |
| W5S_RS11005 | WP_014699936.1 | DNA topoisomerase I |
| W5S_RS13500 | WP_014700412.1 | DNA polymerase III, delta prime subunit |
| W5S_RS17185 | WP_014701083.1 | ssDNA-binding protein |
| W5S_RS01715 | WP_012822131.1 | DNA topoisomerase IV subunit B |
| W5S_RS03370 | WP_005971476.1 | DNA primase |
| W5S_RS15580 | WP_014700794.1 | DNA gyrase subunit A |
| W5S_RS00935 | WP_012822012.1 | primosomal protein N' |
| W5S_RS18425 | WP_014701314.1 | ssDNA-binding protein |
| W5S_RS00020 | WP_014698370.1 | DNA gyrase subunit B |
| W5S_RS16235 | WP_014700900.1 | DNA polymerase III subunit alpha |
| W5S_RS15255 | WP_014700737.1 | DNA polymerase III subunit delta |

| | | |
|---|---|---|
| W5S_RS01770 | WP_014698535.1 | DNA topoisomerase IV subunit A |
| W5S_RS20960 | WP_014701764.1 | DNA-dependent helicase II |
| W5S_RS22185 | WP_014701989.1 | zinc-binding domain of primase-helicase |
| W5S_RS15695 | WP_014700810.1 | DNA polymerase III subunit gamma/tau |
| W5S_RS06990 | WP_014699182.1 | cytochrome bd-I ubiquinol oxidase subunit I |
| W5S_RS14970 | WP_014700693.1 | cytochrome bd-I ubiquinol oxidase subunit I |
| W5S_RS13525 | WP_014700417.1 | beta-ketoacyl-[acyl-carrier-protein synthase II |
| W5S_RS13545 | WP_014700419.1 | 3-oxoacyl-ACP synthase III |
| W5S_RS03410 | WP_012822531.1 | octaprenyl diphosphate synthase |
| W5S_RS06520 | WP_014699103.1 | short-chain dehydrogenase |
| W5S_RS05490 | WP_012822900.1 | pyridoxine 5'-phosphate synthase |
| W5S_RS23135 | WP_014702156.1 | beta-ketoacyl-[acyl-carrier-protein synthase II |
| W5S_RS10235 | WP_005968922.1 | isocitrate dehydrogenase (NADP(+)) |
| W5S_RS13655 | WP_014700440.1 | hypothetical protein |
| W5S_RS03570 | WP_005971563.1 | purine-nucleoside phosphorylase |
| W5S_RS17610 | WP_014701162.1 | phosphatidylserine synthase |
| W5S_RS12745 | WP_014700277.1 | phosphatidylglycerophosphatase B |
| W5S_RS02265 | WP_010284628.1 | cytosol aminopeptidase |
| W5S_RS16315 | WP_005975905.1 | type I methionyl aminopeptidase |
| W5S_RS17335 | WP_014701113.1 | aminoacyl-histidine dipeptidase |
| W5S_RS05715 | WP_014698952.1 | aminopeptidase PepB |
| W5S_RS09765 | WP_014699713.1 | aminopeptidase N |
| W5S_RS22115 | WP_014701975.1 | isoaspartyl peptidase/L-asparaginase |
| W5S_RS06095 | WP_014699026.1 | RNA degradosome polyphosphate kinase |
| W5S_RS05605 | WP_014698935.1 | Pectin acetylesterase pae12A |
| W5S_RS17750 | WP_043899146.1 | sulfate adenylyltransferase |
| W5S_RS17755 | WP_005973508.1 | sulfate adenylyltransferase subunit 2 |
| W5S_RS04995 | WP_012822828.1 | S-ribosylhomocysteine lyase |
| W5S_RS22055 | WP_033072461.1 | phosphopantothenoylcysteine decarboxylase |
| W5S_RS02160 | WP_012822205.1 | aspartate carbamoyltransferase |
| W5S_RS21085 | WP_005969159.1 | adenylosuccinate lyase |
| W5S_RS10210 | WP_014699797.1 | adenylosuccinate lyase |
| W5S_RS14700 | WP_014700648.1 | uridine kinase |
| W5S_RS06175 | WP_014699041.1 | UDP-glucose 6-dehydrogenase |
| W5S_RS17305 | WP_014701108.1 | glutamate 5-kinase |
| W5S_RS06665 | WP_014699128.1 | bifunctional tetrahydrofolate synthase/dihydrofolate synthase |
| W5S_RS16325 | WP_014700913.1 | 2,3,4,5-tetrahydropyridine-2,6-dicarboxylate N-succinyltransferase |
| W5S_RS16445 | WP_014700934.1 | N-acetylglutamate synthase |

| | | |
|---|---|---|
| W5S_RS15660 | WP_014700805.1 | bifunctional UDP-sugar hydrolase/5'-nucleotidase |
| W5S_RS22620 | WP_043898875.1 | bifunctional metallophosphatase/5'-nucleotidase |
| W5S_RS21855 | WP_014701927.1 | phosphoenolpyruvate carboxylase |
| W5S_RS15530 | WP_005976180.1 | 1,4-dihydroxy-2-naphthoyl-CoA synthase |
| W5S_RS22405 | WP_014702034.1 | xylulokinase |
| W5S_RS10685 | WP_014699877.1 | 6-phosphofructokinase |
| W5S_RS00685 | WP_012821982.1 | ATP-dependent 6-phosphofructokinase |
| W5S_RS10840 | WP_015730453.1 | thymidine kinase |
| W5S_RS20905 | WP_014701753.1 | lysophospholipase L2 |
| W5S_RS16135 | WP_014700880.1 | esterase |
| W5S_RS09215 | WP_014699615.1 | metal ABC transporter substrate-binding protein |
| W5S_RS03760 | WP_043898944.1 | methionine ABC transporter substrate-binding protein |
| W5S_RS17665 | WP_014701168.1 | methionine ABC transporter permease |
| W5S_RS17670 | WP_014701169.1 | methionine ABC transporter substrate-binding protein MetQ |
| W5S_RS08250 | WP_014699438.1 | lipoprotein NlpA |
| W5S_RS12165 | WP_014700166.1 | metal ABC transporter substrate-binding protein |
| W5S_RS03755 | WP_012822586.1 | methionine import system permease MetP |
| W5S_RS12175 | WP_014700168.1 | methionine ABC transporter permease |
| W5S_RS06615 | WP_014699120.1 | beta-ketoacyl-[acyl-carrier-protein synthase I |
| W5S_RS14500 | WP_014700604.1 | methylmalonate-semialdehyde dehydrogenase (CoA acylating) |
| W5S_RS15625 | WP_014700799.1 | Cu+ exporting ATPase |
| W5S_RS00440 | WP_014698416.1 | zinc/cadmium/mercury/lead-transporting ATPase |
| W5S_RS09505 | WP_014699663.1 | bifunctional phosphoribosyl-AMP cyclohydrolase/phosphoribosyl-ATP diphosphatase |
| W5S_RS10885 | WP_014699913.1 | cardiolipin synthase A |
| W5S_RS09530 | WP_014699667.1 | histidinol-phosphate transaminase |
| W5S_RS01175 | WP_012822043.1 | uroporphyrinogen decarboxylase |
| W5S_RS22385 | WP_014702030.1 | xylose ABC transporter permease |
| W5S_RS22395 | WP_025919762.1 | D-xylose transporter subunit XylF |
| W5S_RS22390 | WP_014702031.1 | xylose ABC transporter ATP-binding protein |
| W5S_RS00965 | WP_014698476.1 | cystathionine gamma-synthase |
| W5S_RS17985 | WP_014701225.1 | bifunctional dihydroneopterin aldolase/7,8-dihydroneopterin epimerase |
| W5S_RS19595 | WP_014701537.1 | L-ribulose-5-phosphate 4-epimerase |
| W5S_RS12670 | WP_014700264.1 | L-ribulose-5-phosphate 4-epimerase |
| W5S_RS19355 | WP_014701495.1 | 4-hydroxy-3-methylbut-2-enyl diphosphate reductase |
| W5S_RS05435 | WP_014698913.1 | L-aspartate oxidase |
| W5S_RS09330 | WP_014699635.1 | N-acetylmuramoyl-L-alanine amidase |
| W5S_RS19725 | WP_014701561.1 | N-acetylmuramoyl-L-alanine amidase AmiB |

| | | |
|---|---|---|
| W5S_RS16440 | WP_014700933.1 | N-acetylmuramoyl-L-alanine amidase |
| W5S_RS19930 | WP_014701597.1 | CDP-alcohol phosphatidyltransferase |
| W5S_RS11215 | WP_014699975.1 | CDP-alcohol phosphatidyltransferase |
| W5S_RS07175 | WP_014699215.1 | CDP-diacylglycerol--glycerol-3-phosphate 3-phosphatidyltransferase |
| W5S_RS19845 | WP_014701581.1 | fumarate reductase flavoprotein subunit |
| W5S_RS19850 | WP_014701582.1 | succinate dehydrogenase/fumarate reductase iron-sulfur subunit |
| W5S_RS15010 | WP_005973891.1 | succinate dehydrogenase cytochrome b556 small membrane subunit |
| W5S_RS15000 | WP_005973896.1 | succinate dehydrogenase iron-sulfur subunit |
| W5S_RS15005 | WP_005973894.1 | succinate dehydrogenase flavoprotein subunit |
| W5S_RS15015 | WP_014700700.1 | succinate dehydrogenase cytochrome b556 large subunit |
| W5S_RS17990 | WP_014701226.1 | glycerol-3-phosphate acyltransferase |
| W5S_RS13550 | WP_014700420.1 | phosphate acyltransferase |
| W5S_RS03105 | WP_014698703.1 | glycerol-3-phosphate 1-O-acyltransferase |
| W5S_RS19935 | WP_014701598.1 | 1-acyl-sn-glycerol-3-phosphate acyltransferase |
| W5S_RS01775 | WP_012822141.1 | 1-acyl-sn-glycerol-3-phosphate acyltransferase |
| W5S_RS16280 | WP_014700908.1 | phosphatidate cytidylyltransferase |
| W5S_RS19940 | WP_071822931.1 | phosphatidate cytidylyltransferase |
| W5S_RS15915 | WP_014700843.1 | (2E,6E)-farnesyl diphosphate synthase |
| W5S_RS21260 | WP_014701817.1 | dihydroxy-acid dehydratase |
| W5S_RS18640 | WP_014701355.1 | YjhG/YagF family D-xylonate dehydratase |
| W5S_RS12750 | WP_005972518.1 | GTP cyclohydrolase II |
| W5S_RS04460 | WP_012822736.1 | 3,4-dihydroxy-2-butanone-4-phosphate synthase |
| W5S_RS02145 | WP_014698589.1 | anaerobic ribonucleoside triphosphate reductase |
| W5S_RS20890 | WP_014701750.1 | sn-glycerol-3-phosphate dehydrogenase subunit A |
| W5S_RS20810 | WP_014701734.1 | glycerol-3-phosphate dehydrogenase |
| W5S_RS20885 | WP_014701749.1 | anaerobic glycerol-3-phosphate dehydrogenase subunit B |
| W5S_RS17435 | WP_014701131.1 | amidohydrolase |
| W5S_RS04045 | WP_012822655.1 | hydrolase |
| W5S_RS23005 | WP_014702131.1 | urocanate hydratase |
| W5S_RS14900 | WP_014700685.1 | quinolinate synthetase |
| W5S_RS09170 | WP_005967307.1 | S-(hydroxymethyl)glutathione dehydrogenase/class III alcohol dehydrogenase |
| W5S_RS05315 | WP_014698907.1 | 5'-methylthioadenosine/S-adenosylhomocysteine nucleosidase |
| W5S_RS10540 | WP_014699850.1 | L-serine ammonia-lyase |
| W5S_RS10455 | WP_014699835.1 | 5-keto-4-deoxyuronate isomerase |
| W5S_RS20700 | WP_014701714.1 | 5-keto-4-deoxyuronate isomerase |
| W5S_RS01505 | WP_012822094.1 | D-arabinose 5-phosphate isomerase |

| | | |
|---|---|---|
| W5S_RS02960 | WP_012822345.1 | ribose 1,5-bisphosphate phosphokinase PhnN |
| W5S_RS22690 | WP_014702085.1 | guanylate kinase |
| W5S_RS10875 | WP_014699912.1 | oligopeptide ABC transporter ATP-binding protein OppF |
| W5S_RS10870 | WP_014699911.1 | oligopeptide ABC transporter ATP-binding protein OppD |
| W5S_RS08490 | WP_043899008.1 | ABC transporter ATP-binding protein |
| W5S_RS08485 | WP_014699486.1 | ABC transporter ATP-binding protein |
| W5S_RS10865 | WP_014699910.1 | peptide ABC transporter permease |
| W5S_RS20395 | WP_014701655.1 | ABC transporter substrate-binding protein |
| W5S_RS00075 | WP_012821875.1 | ABC transporter substrate-binding protein |
| W5S_RS10855 | WP_014699909.1 | oligopeptide ABC transporter substrate-binding protein OppA |
| W5S_RS08475 | WP_014699484.1 | peptide ABC transporter substrate-binding protein |
| W5S_RS14150 | WP_014700534.1 | ABC transporter substrate-binding protein |
| W5S_RS10850 | WP_014699908.1 | oligopeptide ABC transporter substrate-binding protein OppA |
| W5S_RS09055 | WP_014699585.1 | ABC transporter permease |
| W5S_RS12755 | WP_014700278.1 | murein peptide-binding protein |
| W5S_RS00240 | WP_012821902.1 | ABC transporter substrate-binding protein |
| W5S_RS02610 | WP_012822280.1 | peptide ABC transporter permease |
| W5S_RS09165 | WP_014699606.1 | S-formylglutathione hydrolase |
| W5S_RS16450 | WP_014700935.1 | mandelate racemase |
| W5S_RS16790 | WP_014701001.1 | manganese-dependent inorganic pyrophosphatase |
| W5S_RS18000 | WP_005974741.1 | inorganic pyrophosphatase |
| W5S_RS10730 | WP_014699886.1 | aldehyde dehydrogenase |
| W5S_RS19615 | WP_015731299.1 | erythrose-4-phosphate dehydrogenase |
| W5S_RS22655 | WP_014702079.1 | anion permease |
| W5S_RS12685 | WP_014700266.1 | orotidine-5'-phosphate decarboxylase |
| W5S_RS06240 | WP_014699054.1 | poly(glycerophosphate chain) D-alanine transfer protein |
| W5S_RS05935 | WP_014698997.1 | hydroxyethylthiazole kinase |
| W5S_RS01155 | WP_014698486.1 | thiamine phosphate synthase |
| W5S_RS18250 | WP_043899153.1 | diaminopimelate decarboxylase |
| W5S_RS16605 | WP_014700965.1 | ornithine decarboxylase |
| W5S_RS00975 | WP_012822018.1 | 5,10-methylenetetrahydrofolate reductase |
| W5S_RS21885 | WP_014701933.1 | 5-methyltetrahydropteroyltriglutamate--homocysteine S-methyltransferase |
| W5S_RS06275 | WP_014699060.1 | 5-methyltetrahydropteroyltriglutamate--homocysteine methyltransferase |
| W5S_RS03110 | WP_012822484.1 | diacylglycerol kinase |
| W5S_RS21965 | WP_014701948.1 | lipopolysaccharide heptosyltransferase 1 |
| W5S_RS12905 | WP_014700305.1 | PTS beta-glucoside transporter subunit EIIBCA |

| | | |
|---|---|---|
| W5S_RS21470 | WP_014701850.1 | PTS sugar transporter |
| W5S_RS04425 | WP_012822731.1 | phosphoenolpyruvate--protein phosphotransferase |
| W5S_RS03240 | WP_033071938.1 | PTS beta-glucoside transporter subunit EIIBCA |
| W5S_RS04270 | WP_025920415.1 | PTS beta-glucoside transporter subunit EIIBCA |
| W5S_RS13220 | WP_014700361.1 | PTS beta-glucoside transporter subunit EIIBCA |
| W5S_RS10935 | WP_014699922.1 | tryptophan synthase subunit beta |
| W5S_RS10930 | WP_014699921.1 | tryptophan synthase subunit alpha |
| W5S_RS21395 | WP_014701836.1 | porphobilinogen synthase |
| W5S_RS20770 | WP_014701727.1 | maltodextrin phosphorylase |
| W5S_RS09680 | WP_014699697.1 | tetraacyldisaccharide 4'-kinase |
| W5S_RS08445 | WP_015730244.1 | putrescine aminotransferase |
| W5S_RS11580 | WP_005968483.1 | 3-deoxy-8-phosphooctulonate synthase |
| W5S_RS03655 | WP_012822568.1 | glycine cleavage system protein H |
| W5S_RS03660 | WP_014698750.1 | glycine dehydrogenase (aminomethyl-transferring) |
| W5S_RS00970 | WP_014698477.1 | bifunctional aspartate kinase/homoserine dehydrogenase II |
| W5S_RS19445 | WP_014701509.1 | bifunctional aspartokinase I/homoserine dehydrogenase I |
| W5S_RS19900 | WP_014701591.1 | lysine-sensitive aspartokinase 3 |
| W5S_RS21445 | WP_014701845.1 | acetylglutamate kinase |
| W5S_RS21220 | WP_014701810.1 | sodium:proline symporter |
| W5S_RS12790 | WP_014700286.1 | pyridoxamine 5'-phosphate oxidase |
| W5S_RS13290 | WP_014700372.1 | hypothetical protein |
| W5S_RS01705 | WP_014698529.1 | phosphodiesterase |
| W5S_RS03680 | WP_012822571.1 | phosphodiesterase |
| W5S_RS23030 | WP_014702136.1 | histidine ammonia-lyase |
| W5S_RS14130 | WP_014700530.1 | homocysteine S-methyltransferase |
| W5S_RS21370 | WP_014701832.1 | acetyl-CoA C-acyltransferase FadA |
| W5S_RS06535 | WP_014699106.1 | acetyl-CoA C-acyltransferase FadI |
| W5S_RS20765 | WP_014701726.1 | 4-alpha-glucanotransferase |
| W5S_RS11605 | WP_014700052.1 | glutamyl-tRNA reductase |
| W5S_RS02055 | WP_014698576.1 | type I-E CRISPR-associated protein Cse1/CasA |
| W5S_RS04395 | WP_014698829.1 | cysteine synthase B |
| W5S_RS15150 | WP_014700724.1 | asparagine synthetase B |
| W5S_RS14640 | WP_014700635.1 | pseudaminic acid cytidylyltransferase |
| W5S_RS14635 | WP_081440005.1 | UDP-2,4-diacetamido-2,4,6-trideoxy-beta-L-altropyranose hydrolase |
| W5S_RS14995 | WP_014700699.1 | 2-oxoglutarate dehydrogenase E1 component |
| W5S_RS13370 | WP_043898808.1 | vitamin B12 ABC transporter ATP-binding protein BtuD |
| W5S_RS13380 | WP_014700389.1 | vitamin B12 ABC transporter permease BtuC |
| W5S_RS01005 | WP_012822023.1 | vitamin B12/cobalamin outer membrane transporter |

| | | |
|---|---|---|
| W5S_RS05310 | WP_014698906.1 | cobalamin-binding protein |
| W5S_RS10310 | WP_014699812.1 | glycerophosphodiester phosphodiesterase |
| W5S_RS00605 | WP_014698435.1 | glycerophosphodiester phosphodiesterase |
| W5S_RS19070 | WP_014701439.1 | 2-isopropylmalate synthase |
| W5S_RS15140 | WP_014700722.1 | N-acetylglucosamine-6-phosphate deacetylase |
| W5S_RS01040 | WP_012822026.1 | bifunctional biotin--[acetyl-CoA-carboxylase synthetase/biotin operon repressor |
| W5S_RS01350 | WP_012822067.1 | acetyl-CoA carboxylase biotin carboxyl carrier protein subunit |
| W5S_RS01345 | WP_012822066.1 | acetyl-CoA carboxylase biotin carboxylase subunit |
| W5S_RS15795 | WP_014700825.1 | siderophore biosynthesis protein SbnA |
| W5S_RS16075 | WP_014700868.1 | shikimate kinase II |
| W5S_RS20475 | WP_005969451.1 | shikimate kinase |
| W5S_RS05050 | WP_012822837.1 | bifunctional chorismate mutase/prephenate dehydrogenase |
| W5S_RS17745 | WP_015731092.1 | adenylyl-sulfate kinase |
| W5S_RS20980 | WP_014701768.1 | diaminopimelate epimerase |
| W5S_RS02580 | WP_014698651.1 | rhamnulokinase |
| W5S_RS23045 | WP_014702140.1 | formimidoylglutamate deiminase |
| W5S_RS22700 | WP_005968232.1 | bifunctional GTP diphosphokinase/guanosine-3',5'-bis(diphosphate) 3'-diphosphatase |
| W5S_RS13875 | WP_014700482.1 | DNA starvation/stationary phase protection protein |
| W5S_RS07025 | WP_081483340.1 | nitrite reductase (NAD(P)H) small subunit |
| W5S_RS20410 | WP_014701659.1 | nitrite reductase small subunit |
| W5S_RS07030 | WP_014699190.1 | nitrate reductase |
| W5S_RS03310 | WP_014698729.1 | bifunctional nitrate reductase/sulfite reductase subunit alpha |
| W5S_RS20405 | WP_014701658.1 | nitrite reductase large subunit |
| W5S_RS09150 | WP_014699603.1 | oxidoreductase |
| W5S_RS21095 | WP_014701791.1 | glycerol dehydrogenase |
| W5S_RS08340 | WP_014699454.1 | glycerol dehydrogenase |
| W5S_RS13455 | WP_014700403.1 | Respiratory NADH dehydrogenase 2/cupric reductase |
| W5S_RS17425 | WP_014701129.1 | methylthioribulose 1-phosphate dehydratase |
| W5S_RS15940 | WP_014700847.1 | bifunctional diaminohydroxyphosphoribosylaminopyrimidine deaminase/5-amino-6-(5-phosphoribosylamino)uracil reductase |
| W5S_RS00150 | WP_012821889.1 | alkanesulfonate monooxygenase |
| W5S_RS17385 | WP_014701121.1 | S-methyl-5-thioribose-1-phosphate isomerase |
| W5S_RS21160 | WP_014701800.1 | dTDP-glucose 4,6-dehydratase |
| W5S_RS17930 | WP_014701214.1 | galactarate dehydratase |
| W5S_RS08525 | WP_014699494.1 | CDP-diacylglycerol diphosphatase |

| | | |
|---|---|---|
| W5S_RS21000 | WP_014701774.1 | hydroxymethylbilane synthase |
| W5S_RS06745 | WP_014699141.1 | acetate kinase |
| W5S_RS20655 | WP_014701706.1 | acetate kinase |
| W5S_RS19830 | WP_014701578.1 | phosphatidylserine decarboxylase |
| W5S_RS12525 | WP_014700238.1 | envelope stress response membrane protein PspB |
| W5S_RS17900 | WP_014701207.1 | glycerate kinase |
| W5S_RS10035 | WP_014699763.1 | pyruvate kinase |
| W5S_RS13230 | WP_025918570.1 | pyruvate kinase |
| W5S_RS18235 | WP_014701269.1 | pyruvate kinase |
| W5S_RS12780 | WP_014700284.1 | pyridoxal kinase |
| W5S_RS18930 | WP_014701415.1 | dephospho-CoA kinase |
| W5S_RS03305 | WP_014698728.1 | hypothetical protein |
| W5S_RS13215 | WP_014700360.1 | hypothetical protein |
| W5S_RS00055 | WP_012821871.1 | 6-phospho-beta-glucosidase |
| W5S_RS04265 | WP_014698812.1 | 6-phospho-beta-glucosidase |
| W5S_RS12900 | WP_014700304.1 | 6-phospho-beta-glucosidase |
| W5S_RS11915 | WP_014700117.1 | 6-phospho-beta-glucosidase |
| W5S_RS10450 | WP_005968850.1 | 2-deoxy-D-gluconate 3-dehydrogenase |
| W5S_RS09540 | WP_014699669.1 | ATP phosphoribosyltransferase |
| W5S_RS21235 | WP_005969112.1 | ketol-acid reductoisomerase |
| W5S_RS14610 | WP_014700629.1 | glycosyl transferase |
| W5S_RS01750 | WP_005975568.1 | PTS fructose transporter subunit IIA |
| W5S_RS01745 | WP_010297459.1 | PTS fructose transporter subunit IIB |
| W5S_RS09110 | WP_014699595.1 | PTS fructose transporter subunit IIBC |
| W5S_RS09100 | WP_014699593.1 | bifunctional PTS fructose transporter subunit IIA/HPr protein |
| W5S_RS00585 | WP_014698432.1 | sn-glycerol-3-phosphate ABC transporter substrate-binding protein |
| W5S_RS00600 | WP_014698434.1 | sn-glycerol-3-phosphate import ATP-binding protein UgpC |
| W5S_RS00595 | WP_014698433.1 | sn-glycerol 3-phosphate ABC transporter permease |
| W5S_RS00590 | WP_012821964.1 | glycerol-3-phosphate transporter permease |
| W5S_RS10305 | WP_014699811.1 | ABC transporter substrate-binding protein |
| W5S_RS18925 | WP_014701413.1 | guanosine monophosphate reductase |
| W5S_RS15475 | WP_014700775.1 | spermidine N1-acetyltransferase |
| W5S_RS05665 | WP_014698945.1 | inositol monophosphatase |
| W5S_RS08270 | WP_014699442.1 | amino acid:proton symporter |
| W5S_RS12770 | WP_014700281.1 | amino acid:proton symporter |
| W5S_RS03070 | WP_012822478.1 | C4-dicarboxylate ABC transporter |
| W5S_RS08420 | WP_014699472.1 | anaerobic C4-dicarboxylate transporter |

| | | |
|---|---|---|
| W5S_RS21170 | WP_015731407.1 | UDP-N-acetylglucosamine 2-epimerase (non-hydrolyzing) |
| W5S_RS09770 | WP_014699714.1 | dihydroorotate dehydrogenase (quinone) |
| W5S_RS05755 | WP_014698959.1 | 4-hydroxy-3-methylbut-2-en-1-yl diphosphate synthase |
| W5S_RS18685 | WP_014701363.1 | short-chain dehydrogenase |
| W5S_RS13535 | WP_010276180.1 | beta-ketoacyl-ACP reductase |
| W5S_RS07755 | WP_014699333.1 | short-chain dehydrogenase/reductase SDR |
| W5S_RS21005 | WP_014701775.1 | uroporphyrinogen-III synthase |
| W5S_RS09205 | WP_014699613.1 | glucose sorbosone dehydrogenase |
| W5S_RS19010 | WP_005975233.1 | phospho-N-acetylmuramoyl-pentapeptide-transferase |
| W5S_RS15675 | WP_014700808.1 | adenylate kinase |
| W5S_RS12375 | WP_014700208.1 | sodium-potassium/proton antiporter ChaA |
| W5S_RS20470 | WP_015731352.1 | 3-dehydroquinate synthase |
| W5S_RS20830 | WP_014701738.1 | glucose-1-phosphate adenylyltransferase |
| W5S_RS23055 | WP_014702142.1 | bifunctional N-acetylglucosamine-1-phosphate uridyltransferase/glucosamine-1-phosphate acetyltransferase |
| W5S_RS04305 | WP_012822708.1 | 4-hydroxythreonine-4-phosphate dehydrogenase |
| W5S_RS18650 | WP_005975080.1 | 4-hydroxythreonine-4-phosphate dehydrogenase |
| W5S_RS19295 | WP_014701483.1 | 4-hydroxythreonine-4-phosphate dehydrogenase PdxA |
| W5S_RS10985 | WP_014699932.1 | cob(I)yrinic acid a,c-diamide adenosyltransferase |
| W5S_RS19320 | WP_014701488.1 | dihydrofolate reductase |
| W5S_RS05615 | WP_014698937.1 | serine hydroxymethyltransferase |
| W5S_RS06190 | WP_014699044.1 | bifunctional UDP-glucuronic acid oxidase/UDP-4-amino-4-deoxy-L-arabinose formyltransferase |
| W5S_RS20265 | WP_014701630.1 | serine hydroxymethyltransferase |
| W5S_RS06065 | WP_014699020.1 | galactokinase |
| W5S_RS09105 | WP_014699594.1 | 1-phosphofructokinase |
| W5S_RS21440 | WP_014701844.1 | argininosuccinate lyase |
| W5S_RS11910 | WP_014700116.1 | histidinol-phosphatase |
| W5S_RS09525 | WP_014699666.1 | bifunctional imidazole glycerol-phosphate dehydratase/histidinol phosphatase |
| W5S_RS01045 | WP_012822027.1 | type I pantothenate kinase |
| W5S_RS06165 | WP_014699039.1 | bifunctional methylenetetrahydrofolate dehydrogenase/methenyltetrahydrofolate cyclohydrolase |
| W5S_RS10845 | WP_014699907.1 | bifunctional acetaldehyde-CoA/alcohol dehydrogenase |
| W5S_RS16290 | WP_014700910.1 | 1-deoxy-D-xylulose-5-phosphate reductoisomerase |
| W5S_RS13515 | WP_014700415.1 | aminodeoxychorismate lyase |
| W5S_RS08465 | WP_014699482.1 | AMP nucleosidase |

| | | |
|---|---|---|
| W5S_RS20895 | WP_014701751.1 | MFS transporter |
| W5S_RS15380 | WP_014700759.1 | acetyl-CoA acetyltransferase |
| W5S_RS22070 | WP_014701967.1 | orotate phosphoribosyltransferase |
| W5S_RS06680 | WP_005969887.1 | amidophosphoribosyltransferase |
| W5S_RS09185 | WP_014699609.1 | GTP cyclohydrolase I FolE |
| W5S_RS11565 | WP_014700046.1 | NAD(P) transhydrogenase subunit beta |
| W5S_RS11570 | WP_014700047.1 | NAD(P) transhydrogenase subunit alpha |
| W5S_RS00985 | WP_012822020.1 | NAD(P)(+) transhydrogenase |
| W5S_RS00500 | WP_012821948.1 | branched-chain amino acid ABC transporter permease |
| W5S_RS00490 | WP_025919969.1 | branched chain amino acid ABC transporter substrate-binding protein |
| W5S_RS00510 | WP_005976440.1 | ABC transporter ATP-binding protein |
| W5S_RS00495 | WP_005976436.1 | branched-chain amino acid ABC transporter permease LivH |
| W5S_RS00505 | WP_012821949.1 | ABC transporter ATP-binding protein |
| W5S_RS00885 | WP_012822005.1 | fructose-bisphosphatase class II |
| W5S_RS19675 | WP_014701552.1 | fructose-bisphosphatase class I |
| W5S_RS06135 | WP_014699034.1 | 5-(carboxyamino)imidazole ribonucleotide mutase |
| W5S_RS06130 | WP_014699033.1 | 5-(carboxyamino)imidazole ribonucleotide synthase |
| W5S_RS22415 | WP_014702036.1 | aldehyde dehydrogenase |
| W5S_RS03635 | WP_012822564.1 | alpha-acetolactate decarboxylase |
| W5S_RS22925 | WP_014702122.1 | aspartate--ammonia ligase |
| W5S_RS04195 | WP_033070953.1 | NAD(+) kinase |
| W5S_RS10950 | WP_014699925.1 | glutamine amidotransferase |
| W5S_RS10955 | WP_014699926.1 | anthranilate synthase component I |
| W5S_RS22450 | WP_014702042.1 | PTS mannitol transporter subunit IICBA |
| W5S_RS05860 | WP_014698980.1 | choloylglycine hydrolase |
| W5S_RS16000 | WP_014700855.1 | branched-chain amino acid transport system II carrier protein |
| W5S_RS17370 | WP_014701119.1 | acyl-CoA dehydrogenase |
| W5S_RS21920 | WP_014701939.1 | glycerol-3-phosphate dehydrogenase |
| W5S_RS16500 | WP_014700945.1 | thymidylate synthase |
| W5S_RS06740 | WP_014699140.1 | phosphate acetyltransferase |
| W5S_RS09535 | WP_014699668.1 | histidinol dehydrogenase |
| W5S_RS02790 | WP_012822310.1 | trifunctional nicotinamide-nucleotide adenylyltransferase/ribosylnicotinamide kinase/transcriptional regulator NadR |
| W5S_RS13345 | WP_011093411.1 | idonate transporter |
| W5S_RS18635 | WP_014701354.1 | gluconate permease |
| W5S_RS00340 | WP_043898886.1 | gamma-glutamyltransferase |
| W5S_RS17495 | WP_014701140.1 | gamma-glutamyltransferase |

| | | |
|---|---|---|
| W5S_RS15765 | WP_014700821.1 | ammonium transporter |
| W5S_RS05810 | WP_014698970.1 | GMP synthase (glutamine-hydrolyzing) |
| W5S_RS11125 | WP_014699956.1 | mannose-6-phosphate isomerase |
| W5S_RS22995 | WP_014702129.1 | N-formylglutamate deformylase |
| W5S_RS19895 | WP_014701590.1 | glucose-6-phosphate isomerase |
| W5S_RS22295 | WP_014702012.1 | alpha-hydroxy-acid oxidizing enzyme |
| W5S_RS21750 | WP_014701906.1 | aliphatic sulfonates-binding protein |
| W5S_RS00140 | WP_014698383.1 | aliphatic sulfonates ABC transporter ATP-binding protein |
| W5S_RS17450 | WP_014701134.1 | sulfonate ABC transporter substrate-binding protein |
| W5S_RS21775 | WP_014701911.1 | aliphatic sulfonate ABC transporter ATP-binding protein |
| W5S_RS21770 | WP_014701910.1 | Sulfate ester ABC transporter permease |
| W5S_RS00155 | WP_012821890.1 | aliphatic sulfonate ABC transporter substrate-binding protein |
| W5S_RS08510 | WP_014699491.1 | sulfate ester transporter permease |
| W5S_RS00145 | WP_012821888.1 | sulfonate ABC transporter |
| W5S_RS08505 | WP_014699490.1 | aliphatic sulfonate ABC transporter ATP-binding protein |
| W5S_RS08515 | WP_014699492.1 | aliphatic sulfonates-binding protein |
| W5S_RS20400 | WP_014701656.1 | mandelate racemase |
| W5S_RS21465 | WP_014701849.1 | starvation-sensing protein RspA |
| W5S_RS00125 | WP_012821884.1 | galactonate dehydratase |
| W5S_RS05055 | WP_012822838.1 | bifunctional chorismate mutase/prephenate dehydratase |
| W5S_RS04035 | WP_012822653.1 | O-acetylhomoserine aminocarboxypropyltransferase |
| W5S_RS18045 | WP_014701234.1 | 2',3'-cyclic-nucleotide 2'-phosphodiesterase |
| W5S_RS04055 | WP_014698787.1 | pyruvate:ferredoxin (flavodoxin) oxidoreductase |
| W5S_RS19955 | WP_015731325.1 | malate synthase A |
| W5S_RS17465 | WP_014701137.1 | serine--pyruvate aminotransferase |
| W5S_RS12300 | WP_014700192.1 | ornithine cyclodeaminase |
| W5S_RS06050 | WP_014699017.1 | maltoporin |
| W5S_RS06975 | WP_014699178.1 | PTS glucose transporter subunit IIB |
| W5S_RS02030 | WP_012822182.1 | sucrose porin |
| W5S_RS09990 | WP_014699754.1 | maltoporin |
| W5S_RS20030 | WP_014701612.1 | peptide deformylase |
| W5S_RS14895 | WP_014700684.1 | nicotinamide riboside transporter PnuC |
| W5S_RS10940 | WP_014699923.1 | bifunctional indole-3-glycerol phosphate synthase/phosphoribosylanthranilate isomerase |
| W5S_RS15585 | WP_014700795.1 | bifunctional 3-demethylubiquinone 3-O-methyltransferase/2-octaprenyl-6-hydroxy phenol methylase |

| | | |
|---|---|---|
| W5S_RS01500 | WP_012822093.1 | 3-deoxy-D-manno-octulosonate 8-phosphate phosphatase |
| W5S_RS09760 | WP_014699712.1 | nicotinate phosphoribosyltransferase |
| W5S_RS03275 | WP_012822509.1 | acetylglucosamine-6-sulfatase |
| W5S_RS06970 | WP_014699177.1 | sulfatase |
| W5S_RS17905 | WP_014701209.1 | 2-hydroxy-3-oxopropionate reductase |
| W5S_RS20615 | WP_014701700.1 | aquaporin |
| W5S_RS00895 | WP_014698472.1 | aquaporin |
| W5S_RS15410 | WP_014700764.1 | phosphoribosylaminoimidazolesuccinocarboxamide synthase |
| W5S_RS15460 | WP_014700772.1 | phosphoribosylformylglycinamidine cyclo-ligase |
| W5S_RS15455 | WP_081483327.1 | uracil phosphoribosyltransferase |
| W5S_RS21215 | WP_014701809.1 | trifunctional transcriptional regulator/proline dehydrogenase/L-glutamate gamma-semialdehyde dehydrogenase |
| W5S_RS21345 | WP_014701828.1 | protoporphyrinogen oxidase |
| W5S_RS22080 | WP_014701969.1 | dipeptide epimerase |
| W5S_RS06345 | WP_014699070.1 | dipeptide epimerase |
| W5S_RS00170 | WP_014698385.1 | beta-glucosidase |
| W5S_RS17780 | WP_014701185.1 | hypothetical protein |
| W5S_RS17910 | WP_014701210.1 | 5-keto-4-deoxy-D-glucarate aldolase |
| W5S_RS19950 | WP_014701601.1 | isocitrate lyase |
| W5S_RS13585 | WP_014700425.1 | dihydroorotase |
| W5S_RS23000 | WP_014702130.1 | imidazolonepropionase |
| W5S_RS01160 | WP_014698487.1 | phosphomethylpyrimidine synthase ThiC |
| W5S_RS06100 | WP_014699027.1 | phosphate ABC transporter permease |
| W5S_RS16010 | WP_014700857.1 | phosphate ABC transporter substrate-binding protein |
| W5S_RS23160 | WP_014702161.1 | phosphate ABC transporter substrate-binding protein |
| W5S_RS23165 | WP_014702162.1 | phosphate transporter permease subunit PstC |
| W5S_RS23170 | WP_014702163.1 | phosphate transporter permease subunit PtsA |
| W5S_RS23175 | WP_014702164.1 | phosphate ABC transporter ATP-binding protein |
| W5S_RS06110 | WP_014699029.1 | phosphate ABC transporter ATP-binding protein |
| W5S_RS06105 | WP_014699028.1 | phosphate ABC transporter, permease protein PstA |
| W5S_RS14990 | WP_014700698.1 | dihydrolipoamide succinyltransferase |
| W5S_RS21200 | WP_014701807.1 | exopolyphosphatase |
| W5S_RS03390 | WP_012822528.1 | malate dehydrogenase |
| W5S_RS10115 | WP_014699777.1 | malate/lactate/ureidoglycolate dehydrogenase |
| W5S_RS08360 | WP_014699458.1 | malate/lactate/ureidoglycolate dehydrogenase |
| W5S_RS12280 | WP_014700189.1 | Bme24 |
| W5S_RS14830 | WP_014700672.1 | UDP-glucose 4-epimerase |

| | | |
|---|---|---|
| W5S_RS05395 | WP_014698911.1 | glutamate--tRNA ligase |
| W5S_RS20340 | WP_014701644.1 | aspartate aminotransferase family protein |
| W5S_RS04640 | WP_012822770.1 | cystathionine beta-lyase |
| W5S_RS01980 | WP_012822174.1 | cystathionine beta-lyase |
| W5S_RS01545 | WP_012822102.1 | UDP-N-acetylglucosamine 1-carboxyvinyltransferase |
| W5S_RS19375 | WP_014701499.1 | bifunctional riboflavin kinase/FMN adenylyltransferase |
| W5S_RS04310 | WP_014698818.1 | triose-phosphate isomerase |
| W5S_RS00865 | WP_014698470.1 | triose-phosphate isomerase |
| W5S_RS19605 | WP_014701539.1 | class II fructose-bisphosphate aldolase |
| W5S_RS14070 | WP_014700518.1 | isopentenyl-diphosphate Delta-isomerase |
| W5S_RS09400 | WP_014699646.1 | thiol reductant ABC exporter subunit CydD |
| W5S_RS06985 | WP_014699181.1 | cytochrome d ubiquinol oxidase subunit II |
| W5S_RS15865 | WP_014700837.1 | cytochrome ubiquinol oxidase subunit I |
| W5S_RS15870 | WP_014700838.1 | cytochrome o ubiquinol oxidase subunit III |
| W5S_RS15860 | WP_014700836.1 | cytochrome ubiquinol oxidase subunit II |
| W5S_RS09395 | WP_014699645.1 | amino acid ABC transporter ATP-binding protein |
| W5S_RS14965 | WP_014700692.1 | cytochrome d ubiquinol oxidase subunit II |
| W5S_RS15875 | WP_011092734.1 | cytochrome o ubiquinol oxidase subunit IV |
| W5S_RS22745 | WP_014702094.1 | glutamine synthetase |
| W5S_RS06660 | WP_005969895.1 | acetyl-CoA carboxylase carboxyl transferase subunit beta |
| W5S_RS16230 | WP_014700899.1 | acetyl-CoA carboxylase carboxyl transferase subunit alpha |
| W5S_RS00400 | WP_012821933.1 | C4-dicarboxylate transporter DctA |
| W5S_RS17790 | WP_014701187.1 | phosphoadenosine phosphosulfate reductase |
| W5S_RS22025 | WP_014701962.1 | pantetheine-phosphate adenylyltransferase |
| W5S_RS14075 | WP_033071719.1 | beta-glucosidase |
| W5S_RS03975 | WP_014698781.1 | 4-hydroxy-2-oxoglutarate aldolase / 2-dehydro-3-deoxyphosphogluconate aldolase |
| W5S_RS10050 | WP_005967771.1 | ketohydroxyglutarate aldolase |
| W5S_RS14570 | WP_014700618.1 | glucose-1-phosphate thymidylyltransferase |
| W5S_RS06010 | WP_014699011.1 | D-lactate dehydrogenase |
| W5S_RS12450 | WP_014700224.1 | lactate dehydrogenase |
| W5S_RS22490 | WP_014702049.1 | bifunctional glyoxylate/hydroxypyruvate reductase B |
| W5S_RS09135 | WP_025918944.1 | glycerate dehydrogenase |
| W5S_RS21265 | WP_014701818.1 | branched chain amino acid aminotransferase |
| W5S_RS10015 | WP_014699759.1 | zinc ABC transporter ATP-binding protein ZnuC |
| W5S_RS10010 | WP_014699758.1 | zinc ABC transporter permease |
| W5S_RS18515 | WP_043899159.1 | ABC transporter |
| W5S_RS18520 | WP_014701332.1 | iron ABC transporter |

| | | |
|---|---|---|
| W5S_RS11860 | WP_014700105.1 | manganese ABC transporter ATP-binding protein |
| W5S_RS11865 | WP_014700106.1 | Manganese transport system membrane protein MntB |
| W5S_RS18525 | WP_014701333.1 | metal ABC transporter substrate-binding protein |
| W5S_RS10020 | WP_014699760.1 | zinc ABC transporter substrate-binding protein |
| W5S_RS17455 | WP_014701135.1 | aspartate aminotransferase family protein |
| W5S_RS02785 | WP_043898935.1 | 5-formyltetrahydrofolate cyclo-ligase |
| W5S_RS14300 | WP_014700564.1 | 8-amino-7-oxononanoate synthase |
| W5S_RS23060 | WP_014702143.1 | glutamine--fructose-6-phosphate aminotransferase |
| W5S_RS06775 | WP_014699146.1 | aminotransferase AlaT |
| W5S_RS05905 | WP_014698990.1 | alcohol dehydrogenase |
| W5S_RS18660 | WP_014701358.1 | alcohol dehydrogenase |
| W5S_RS05895 | WP_014698987.1 | zinc-dependent alcohol dehydrogenase |
| W5S_RS15935 | WP_014700846.1 | 6,7-dimethyl-8-ribityllumazine synthase |
| W5S_RS22400 | WP_014702033.1 | xylose isomerase |
| W5S_RS17915 | WP_014701211.1 | glucarate dehydratase |
| W5S_RS13300 | WP_014700374.1 | phosphoenolpyruvate synthase |
| W5S_RS03565 | WP_012822553.1 | phosphopentomutase |
| W5S_RS06340 | WP_014699069.1 | D-amino acid aminotransferase |
| W5S_RS21255 | WP_014701816.1 | PLP-dependent threonine dehydratase |
| W5S_RS23915 | WP_014699501.1 | serine/threonine dehydratase |
| W5S_RS14535 | WP_014700611.1 | chorismate mutase |
| W5S_RS10830 | WP_005968726.1 | UTP--glucose-1-phosphate uridylyltransferase |
| W5S_RS14565 | WP_014700617.1 | UTP--glucose-1-phosphate uridylyltransferase |
| W5S_RS06525 | WP_014699104.1 | long-chain fatty acid transporter |
| W5S_RS05295 | WP_014698904.1 | aspartate aminotransferase family protein |
| W5S_RS19510 | WP_025920194.1 | ribose 5-phosphate isomerase A |
| W5S_RS04285 | WP_014698816.1 | ribose 5-phosphate isomerase B |
| W5S_RS03365 | WP_014698735.1 | RNA polymerase sigma factor RpoD |
| W5S_RS01115 | WP_012822031.1 | DNA-directed RNA polymerase subunit beta' |
| W5S_RS13155 | WP_014700349.1 | heme lysase NrfEFG subunit NrfG |
| W5S_RS06610 | WP_014699119.1 | bifunctional tRNA (5-methylaminomethyl-2-thiouridine)(34)-methyltransferase MnmD/FAD-dependent 5-carboxymethylaminomethyl-2-thiouridine(34) oxidoreductase MnmC |
| W5S_RS05475 | WP_012822897.1 | ribonuclease III |
| W5S_RS03490 | WP_012822541.1 | transcription termination protein NusA |
| W5S_RS17975 | WP_014701223.1 | multifunctional CCA tRNA nucleotidyl transferase/2'3'-cyclic phosphodiesterase/2'nucleotidase/phosphatase |
| W5S_RS05020 | WP_012822832.1 | tRNA (guanosine(37)-N1)-methyltransferase TrmD |

| | | |
|---|---|---|
| W5S_RS20065 | WP_005970247.1 | DNA-directed RNA polymerase subunit alpha |
| W5S_RS01110 | WP_012822030.1 | DNA-directed RNA polymerase subunit beta |
| W5S_RS00620 | WP_012821970.1 | tRNA (uridine(34)/cytosine(34)/5-carboxymethylaminomethyluridine(34)-2'-O)-methyltransferase TrmL |
| W5S_RS20005 | WP_014701607.1 | shikimate dehydrogenase |
| W5S_RS13375 | WP_014700388.1 | glutathione peroxidase |
| W5S_RS02025 | WP_014698572.1 | PTS sucrose IIBC component |
| W5S_RS13195 | WP_014700354.1 | Pyrimidine-specific ribonucleoside hydrolase rihA |
| W5S_RS00090 | WP_014698378.1 | Pyrimidine ribonucleoside hydrolase rihA |
| W5S_RS00085 | WP_014698377.1 | Pyrimidine ribonucleoside hydrolase rihA |
| W5S_RS18710 | WP_014701368.1 | ribokinase |
| W5S_RS22870 | WP_014702113.1 | ribokinase |
| W5S_RS20785 | WP_014701729.1 | ribokinase |
| W5S_RS15700 | WP_014700811.1 | adenine phosphoribosyltransferase |
| W5S_RS10945 | WP_014699924.1 | anthranilate phosphoribosyltransferase |
| W5S_RS15525 | WP_014700785.1 | o-succinylbenzoate synthase |
| W5S_RS17710 | WP_005973491.1 | 2-C-methyl-D-erythritol 2,4-cyclodiphosphate synthase |
| W5S_RS19565 | WP_014701531.1 | 3-dehydro-L-gulonate 2-dehydrogenase |
| W5S_RS17655 | WP_014701166.1 | D-glycero-beta-D-manno-heptose 1,7-bisphosphate 7-phosphatase |
| W5S_RS09515 | WP_014699664.1 | 1-(5-phosphoribosyl)-5-((5-phosphoribosylamino)methylideneamino)imidazole-4-carboxamide isomerase |
| W5S_RS03560 | WP_014698743.1 | thymidine phosphorylase |
| W5S_RS01590 | WP_012822108.1 | acetoin reductase |
| W5S_RS15980 | WP_005975991.1 | ACP phosphodiesterase |
| W5S_RS14980 | WP_014700696.1 | succinate--CoA ligase subunit alpha |
| W5S_RS14985 | WP_014700697.1 | succinate--CoA ligase subunit beta |
| W5S_RS17445 | WP_014701133.1 | glycyl radical enzyme |
| W5S_RS18690 | WP_014701364.1 | glycyl radical enzyme |
| W5S_RS09465 | WP_005967554.1 | formate acetyltransferase |
| W5S_RS05420 | WP_005973551.1 | autonomous glycyl radical cofactor GrcA |
| W5S_RS20625 | WP_014701701.1 | glycyl radical enzyme |
| W5S_RS02805 | WP_014698672.1 | protein smp |
| W5S_RS06925 | WP_014699169.1 | hypothetical protein |
| W5S_RS20805 | WP_014701733.1 | thiosulfate sulfurtransferase |
| W5S_RS13360 | WP_043898807.1 | phosphogluconate dehydratase |
| W5S_RS14655 | WP_014700639.1 | undecaprenyl-phosphate galactose phosphotransferase WbaP |
| W5S_RS08355 | WP_014699457.1 | aliphatic sulfonates-binding protein |

| | | |
|---|---|---|
| W5S_RS08370 | WP_015730232.1 | taurine ABC transporter permease |
| W5S_RS12240 | WP_014700180.1 | aspartate aminotransferase family protein |
| W5S_RS15090 | WP_014700713.1 | phosphoglucomutase, alpha-D-glucose phosphate-specific |
| W5S_RS05220 | WP_014698896.1 | 3-methyl-2-oxobutanoate hydroxymethyltransferase |
| W5S_RS04295 | WP_014698817.1 | 2-keto-3-deoxygluconate permease |
| W5S_RS20695 | WP_014701713.1 | 2-keto-3-deoxygluconate transporter |
| W5S_RS20825 | WP_014701737.1 | starch synthase |
| W5S_RS20605 | WP_014701698.1 | methionine adenosyltransferase |
| W5S_RS19640 | WP_014701545.1 | S-adenosylmethionine synthase |
| W5S_RS21980 | WP_014701952.1 | lipopolysaccharide core heptosyltransferase RfaQ |
| W5S_RS07145 | WP_014699211.1 | D-serine/D-alanine/glycine transporter |
| W5S_RS00545 | WP_014698424.1 | hydroxypyruvate isomerase |
| W5S_RS04715 | WP_081483311.1 | spermidine/putrescine ABC transporter permease |
| W5S_RS10170 | WP_014699789.1 | spermidine/putrescine ABC transporter substrate-binding protein PotD |
| W5S_RS09270 | WP_005967349.1 | polyamine ABC transporter ATP-binding protein |
| W5S_RS09275 | WP_014699625.1 | putrescine ABC transporter permease |
| W5S_RS14440 | WP_014700592.1 | ABC transporter |
| W5S_RS09280 | WP_014699626.1 | putrescine ABC transporter permease PotI |
| W5S_RS12250 | WP_014700182.1 | spermidine/putrescine ABC transporter ATP-binding protein |
| W5S_RS10185 | WP_014699792.1 | putrescine/spermidine ABC transporter ATP-binding protein |
| W5S_RS12220 | WP_014700177.1 | polyamine ABC transporter substrate-binding protein |
| W5S_RS10175 | WP_014699790.1 | spermidine/putrescine ABC transporter permease PotC |
| W5S_RS12225 | WP_014700178.1 | spermidine/putrescine ABC transporter substrate-binding protein |
| W5S_RS12215 | WP_014700176.1 | polyamine ABC transporter permease |
| W5S_RS10180 | WP_014699791.1 | spermidine/putrescine ABC transporter permease PotB |
| W5S_RS21985 | WP_014701953.1 | glucosyltransferase |
| W5S_RS03100 | WP_012822482.1 | 4-hydroxybenzoate octaprenyltransferase |
| W5S_RS22425 | WP_014702038.1 | superoxide dismutase [Mn |
| W5S_RS18870 | WP_014701402.1 | pyruvate dehydrogenase complex dihydrolipoyllysine-residue acetyltransferase |
| W5S_RS02115 | WP_014698586.1 | aspartate aminotransferase family protein |
| W5S_RS19635 | WP_014701544.1 | arginine decarboxylase |
| W5S_RS22795 | WP_014702103.1 | oxygen-independent coproporphyrinogen III oxidase |
| W5S_RS06125 | WP_014699032.1 | Threonine/serine transporter tdcC |

| | | |
|---|---|---|
| W5S_RS14770 | WP_014700660.1 | glucokinase |
| W5S_RS19425 | WP_005973198.1 | transaldolase |
| W5S_RS04280 | WP_014698815.1 | transaldolase |
| W5S_RS18040 | WP_014701233.1 | 3'(2'),5'-bisphosphate nucleotidase CysQ |
| W5S_RS01300 | WP_014698499.1 | sodium:pantothenate symporter |
| W5S_RS03285 | WP_014698723.1 | beta-phosphoglucomutase |
| W5S_RS00305 | WP_014698397.1 | valine--pyruvate transaminase |
| W5S_RS05940 | WP_014698998.1 | hydroxymethylpyrimidine/phosphomethylpyrimidine kinase |
| W5S_RS12030 | WP_014700140.1 | MFS transporter |
| W5S_RS16245 | WP_014700902.1 | lipid-A-disaccharide synthase |
| W5S_RS21165 | WP_014701801.1 | UDP-N-acetyl-D-mannosamine dehydrogenase |
| W5S_RS05265 | WP_025920215.1 | iron-hydroxamate transporter ATP-binding subunit |
| W5S_RS05275 | WP_014698902.1 | Fe3+-hydroxamate ABC transporter permease FhuB |
| W5S_RS05270 | WP_012822869.1 | iron-hydroxamate transporter substrate-binding subunit |
| W5S_RS22300 | WP_014702013.1 | aldehyde dehydrogenase |
| W5S_RS15535 | WP_081483346.1 | 2-succinyl-6-hydroxy-2,4-cyclohexadiene-1-carboxylate synthase |
| W5S_RS19440 | WP_005973192.1 | homoserine kinase |
| W5S_RS19885 | WP_014701588.1 | EF-P beta-lysylation protein EpmB |
| W5S_RS08335 | WP_014699453.1 | glutamate/aspartate:proton symporter GltP |
| W5S_RS11615 | WP_014700054.1 | 4-(cytidine 5'-diphospho)-2-C-methyl-D-erythritol kinase |
| W5S_RS19610 | WP_005973111.1 | phosphoglycerate kinase |
| W5S_RS19685 | WP_014701554.1 | adenylosuccinate synthase |
| W5S_RS15880 | WP_005976015.1 | protoheme IX farnesyltransferase |
| W5S_RS05195 | WP_005969011.1 | carbonate dehydratase |
| W5S_RS01290 | WP_012822056.1 | carbonic anhydrase |
| W5S_RS20035 | WP_014701613.1 | methionyl-tRNA formyltransferase |
| W5S_RS13750 | WP_005970602.1 | methylglyoxal synthase |
| W5S_RS05320 | WP_014698908.1 | dGTPase |
| W5S_RS03180 | WP_014698706.1 | uronate isomerase |
| W5S_RS06585 | WP_005969914.1 | chorismate synthase |
| W5S_RS19275 | WP_014701479.1 | bifunctional tRNA pseudouridine(32) synthase/ribosomal large subunit pseudouridine synthase RluA |
| W5S_RS10980 | WP_014699931.1 | 23S rRNA pseudouridylate synthase B |
| W5S_RS03505 | WP_014698738.1 | tRNA pseudouridine(55) synthase |
| W5S_RS06645 | WP_014699126.1 | tRNA pseudouridine(38-40) synthase |
| W5S_RS13570 | WP_043898811.1 | 23S rRNA pseudouridine(955/2504/2580) synthase |
| W5S_RS09035 | WP_014699581.1 | 16S rRNA pseudouridine(516) synthase |

| | | |
|---|---|---|
| W5S_RS10230 | WP_014699800.1 | 23S rRNA pseudouridine(2457) synthase |
| W5S_RS05070 | WP_012822839.1 | 23S rRNA pseudouridine(1911/1915/1917) synthase |
| W5S_RS15130 | WP_014700720.1 | PTS N-acetylglucosamine IIBC component |
| W5S_RS14725 | WP_014700653.1 | formate dehydrogenase cytochrome b556 subunit |
| W5S_RS08275 | WP_014699443.1 | formate dehydrogenase subunit alpha |
| W5S_RS14720 | WP_014700652.1 | formate dehydrogenase subunit beta |
| W5S_RS14715 | WP_014700651.1 | formate dehydrogenase-N subunit alpha |
| W5S_RS09345 | WP_014699638.1 | hydroxylamine reductase |
| W5S_RS09745 | WP_014699709.1 | aspartate aminotransferase |
| W5S_RS18350 | WP_014701294.1 | aromatic amino acid aminotransferase |
| W5S_RS19310 | WP_014701486.1 | bis(5'-nucleosyl)-tetraphosphatase (symmetrical) |
| W5S_RS02765 | WP_014698669.1 | 2-octaprenyl-6-methoxyphenyl hydroxylase |
| W5S_RS14625 | WP_014700632.1 | pseudaminic acid synthase |
| W5S_RS03095 | WP_014698702.1 | chorismate lyase |
| W5S_RS20160 | WP_005970274.1 | 30S ribosomal protein S3 |
| W5S_RS22465 | WP_005968098.1 | glycine--tRNA ligase subunit alpha |
| W5S_RS20180 | WP_005970277.1 | 50S ribosomal protein L23 |
| W5S_RS03375 | WP_001144069.1 | 30S ribosomal protein S21 |
| W5S_RS05760 | WP_014698960.1 | histidine--tRNA ligase |
| W5S_RS20185 | WP_005970278.1 | 50S ribosomal protein L4 |
| W5S_RS06155 | WP_014699037.1 | cysteine--tRNA ligase |
| W5S_RS20190 | WP_014701620.1 | 50S ribosomal protein L3 |
| W5S_RS15735 | WP_014700818.1 | 50S ribosomal protein L31 |
| W5S_RS00940 | WP_005973342.1 | 50S ribosomal protein L31 |
| W5S_RS12785 | WP_014700285.1 | tyrosine--tRNA ligase |
| W5S_RS20165 | WP_005970275.1 | 50S ribosomal protein L22 |
| W5S_RS19875 | WP_014701586.1 | entericidin B |
| W5S_RS20120 | WP_005970261.1 | 30S ribosomal protein S8 |
| W5S_RS20170 | WP_004929772.1 | 30S ribosomal protein S19 |
| W5S_RS20220 | WP_005969574.1 | 30S ribosomal protein S7 |
| W5S_RS20175 | WP_014701619.1 | 50S ribosomal protein L2 |
| W5S_RS03790 | WP_014698763.1 | lysine--tRNA ligase |
| W5S_RS20140 | WP_000613954.1 | 50S ribosomal protein L14 |
| W5S_RS14710 | WP_014700650.1 | methionine--tRNA ligase |
| W5S_RS20135 | WP_005970267.1 | 50S ribosomal protein L24 |
| W5S_RS20145 | WP_005970270.1 | 30S ribosomal protein S17 |
| W5S_RS05025 | WP_005970359.1 | 50S ribosomal protein L19 |
| W5S_RS03510 | WP_005971532.1 | 30S ribosomal protein S15 |
| W5S_RS20150 | WP_005970272.1 | 50S ribosomal protein L29 |

| | | |
|---|---|---|
| W5S_RS20225 | WP_005969567.1 | 30S ribosomal protein S12 |
| W5S_RS20155 | WP_005970273.1 | 50S ribosomal protein L16 |
| W5S_RS01575 | WP_005975490.1 | 30S ribosomal protein S9 |
| W5S_RS03420 | WP_005971498.1 | 50S ribosomal protein L27 |
| W5S_RS20070 | WP_005970249.1 | 30S ribosomal protein S4 |
| W5S_RS04940 | WP_014698885.1 | alanine--tRNA ligase |
| W5S_RS20110 | WP_005970258.1 | 50S ribosomal protein L18 |
| W5S_RS10360 | WP_005968885.1 | phenylalanine--tRNA ligase subunit alpha |
| W5S_RS01570 | WP_005975488.1 | 50S ribosomal protein L13 |
| W5S_RS18080 | WP_014701242.1 | 50S ribosomal protein L9 |
| W5S_RS15245 | WP_014700735.1 | leucine--tRNA ligase |
| W5S_RS18085 | WP_005974788.1 | 30S ribosomal protein S18 |
| W5S_RS05010 | WP_005970362.1 | 30S ribosomal protein S16 |
| W5S_RS02255 | WP_014698600.1 | valine--tRNA ligase |
| W5S_RS01100 | WP_005970333.1 | 50S ribosomal protein L10 |
| W5S_RS20085 | WP_002227352.1 | 50S ribosomal protein L36 |
| W5S_RS03415 | WP_005971496.1 | 50S ribosomal protein L21 |
| W5S_RS01095 | WP_012822029.1 | 50S ribosomal protein L1 |
| W5S_RS22045 | WP_005967968.1 | 50S ribosomal protein L28 |
| W5S_RS22460 | WP_014702044.1 | glycine--tRNA ligase subunit beta |
| W5S_RS19380 | WP_005973217.1 | 30S ribosomal protein S20 |
| W5S_RS13555 | WP_005970518.1 | 50S ribosomal protein L32 |
| W5S_RS20080 | WP_005970252.1 | 30S ribosomal protein S13 |
| W5S_RS09960 | WP_014699748.1 | aspartate--tRNA ligase |
| W5S_RS20125 | WP_010286114.1 | 30S ribosomal protein S14 |
| W5S_RS09885 | WP_014699735.1 | arginine--tRNA ligase |
| W5S_RS20095 | WP_005970256.1 | 50S ribosomal protein L15 |
| W5S_RS20195 | WP_001181005.1 | 30S ribosomal protein S10 |
| W5S_RS18095 | WP_005974793.1 | 30S ribosomal protein S6 |
| W5S_RS09430 | WP_014699651.1 | serine--tRNA ligase |
| W5S_RS20105 | WP_005970257.1 | 30S ribosomal protein S5 |
| W5S_RS20445 | WP_014701665.1 | tryptophan--tRNA ligase |
| W5S_RS23325 | WP_005976668.1 | 50S ribosomal protein L34 |
| W5S_RS10350 | WP_005968897.1 | 50S ribosomal protein L35 |
| W5S_RS22040 | WP_002208990.1 | 50S ribosomal protein L33 |
| W5S_RS01105 | WP_014698481.1 | 50S ribosomal protein L7/L12 |
| W5S_RS10340 | WP_014699817.1 | threonine--tRNA ligase |
| W5S_RS16220 | WP_043898821.1 | tRNA(Ile)-lysidine synthetase |
| W5S_RS09755 | WP_014699711.1 | asparagine--tRNA ligase |

| | | |
|---|---|---|
| W5S_RS20115 | WP_005970259.1 | 50S ribosomal protein L6 |
| W5S_RS10365 | WP_014699819.1 | phenylalanine--tRNA ligase subunit beta |
| W5S_RS24150 | WP_005976091.1 | 50S ribosomal protein L36 |
| W5S_RS16310 | WP_010284816.1 | 30S ribosomal protein S2 |
| W5S_RS20130 | WP_005970265.1 | 50S ribosomal protein L5 |
| W5S_RS10355 | WP_005968887.1 | 50S ribosomal protein L20 |
| W5S_RS19370 | WP_014701498.1 | isoleucine--tRNA ligase |
| W5S_RS20060 | WP_005970246.1 | 50S ribosomal protein L17 |
| W5S_RS19560 | WP_014701530.1 | gluconolactonase |
| W5S_RS00115 | WP_014698381.1 | 2-dehydro-3-deoxygalactonokinase |
| W5S_RS12610 | WP_014700252.1 | flavocytochrome c |
| W5S_RS19860 | WP_014701584.1 | fumarate reductase subunit D |
| W5S_RS04520 | WP_025919880.1 | fumarate reductase |
| W5S_RS04515 | WP_014698838.1 | flavocytochrome c |
| W5S_RS19855 | WP_025920292.1 | fumarate reductase subunit C |
| W5S_RS21150 | WP_014701798.1 | dTDP-4-amino-4,6-dideoxy-D-glucose transaminase |
| W5S_RS19020 | WP_014701431.1 | UDP-N-acetylmuramoyl-L-alanyl-D-glutamate--2,6-diaminopimelate ligase |
| W5S_RS12840 | WP_014700293.1 | cyclopropane-fatty-acyl-phospholipid synthase |
| W5S_RS03175 | WP_014698705.1 | MFS transporter |
| W5S_RS12620 | WP_014700254.1 | MFS transporter |
| W5S_RS21155 | WP_014701799.1 | TDP-D-fucosamine acetyltransferase |
| W5S_RS22715 | WP_014702088.1 | purine permease yicE |
| W5S_RS20455 | WP_014701667.1 | ribulose-phosphate 3-epimerase |
| W5S_RS06810 | WP_014699151.1 | NADH-quinone oxidoreductase subunit G |
| W5S_RS06815 | WP_011094550.1 | NADH-quinone oxidoreductase subunit H |
| W5S_RS06805 | WP_014699150.1 | NADH-quinone oxidoreductase subunit F |
| W5S_RS06795 | WP_014699149.1 | NADH-quinone oxidoreductase subunit C/D |
| W5S_RS06800 | WP_005969842.1 | NADH-quinone oxidoreductase subunit E |
| W5S_RS06790 | WP_014699148.1 | NADH dehydrogenase subunit B |
| W5S_RS06825 | WP_005969837.1 | NADH:ubiquinone oxidoreductase subunit J |
| W5S_RS06830 | WP_005969835.1 | NADH-quinone oxidoreductase subunit K |
| W5S_RS06840 | WP_014699153.1 | NADH-quinone oxidoreductase subunit M |
| W5S_RS06820 | WP_010278884.1 | NADH-quinone oxidoreductase subunit I |
| W5S_RS06845 | WP_014699154.1 | NADH-quinone oxidoreductase subunit N |
| W5S_RS06785 | WP_005969847.1 | NADH-quinone oxidoreductase subunit A |
| W5S_RS06835 | WP_014699152.1 | NADH-quinone oxidoreductase subunit L |
| W5S_RS16320 | WP_014700912.1 | bifunctional uridylyltransferase/uridylyl-removing protein |
| W5S_RS22445 | WP_014702041.1 | mannitol-1-phosphate 5-dehydrogenase |

| | | |
|---|---|---|
| W5S_RS05225 | WP_014698897.1 | 2-amino-4-hydroxy-6-hydroxymethyldihydropteridine diphosphokinase |
| W5S_RS09520 | WP_014699665.1 | imidazole glycerol phosphate synthase subunit HisH |
| W5S_RS09510 | WP_005967575.1 | imidazole glycerol phosphate synthase cyclase subunit |
| W5S_RS14455 | WP_014700595.1 | 5-deoxy-glucuronate isomerase |
| W5S_RS17330 | WP_005975741.1 | xanthine phosphoribosyltransferase |
| W5S_RS15375 | WP_014700758.1 | succinyl-CoA--3-ketoacid-CoA transferase |
| W5S_RS15370 | WP_014700757.1 | acetate CoA-transferase subunit alpha |
| W5S_RS05620 | WP_014698938.1 | 3-phenylpropionic acid transporter |
| W5S_RS06935 | WP_014699171.1 | glutamine amidotransferase |
| W5S_RS16285 | WP_014700909.1 | (2E,6E)-farnesyl- diphosphate-specific ditrans,polycis-undecaprenyl-diphosphate synthase |
| W5S_RS00410 | WP_012821935.1 | ketodeoxygluconokinase |
| W5S_RS06855 | WP_014699155.1 | glucohydrolase |
| W5S_RS05910 | WP_014698991.1 | 3-isopropylmalate dehydratase large subunit |
| W5S_RS19080 | WP_014701441.1 | 3-isopropylmalate dehydratase large subunit |
| W5S_RS19085 | WP_014701442.1 | 3-isopropylmalate dehydratase small subunit |
| W5S_RS05915 | WP_025919148.1 | 3-isopropylmalate dehydratase small subunit |
| W5S_RS11080 | WP_005968627.1 | L-arabinose ABC transporter permease AraH |
| W5S_RS11070 | WP_005968631.1 | arabinose ABC transporter substrate-binding protein |
| W5S_RS11075 | WP_014699947.1 | Arabinose import ATP-binding protein AraG |
| W5S_RS19075 | WP_014701440.1 | 3-isopropylmalate dehydrogenase |
| W5S_RS04245 | WP_014698809.1 | ATP-binding protein of ABC mannitol transporter |
| W5S_RS06045 | WP_014699016.1 | Maltose/maltodextrin ABC transporter ATP-binding protein MalK |
| W5S_RS06035 | WP_043898762.1 | arabinogalactan ABC transporter permease |
| W5S_RS06040 | WP_014699015.1 | Maltosaccharide ABC transporter maltosaccharide-binding protein |
| W5S_RS06030 | WP_011094699.1 | arabinogalactan ABC transporter permease |
| W5S_RS14190 | WP_014700543.1 | ferrichrome ABC transporter permease |
| W5S_RS09660 | WP_014699694.1 | iron ABC transporter ATP-binding protein |
| W5S_RS05655 | WP_005970186.1 | iron-siderophore ABC transporter permease |
| W5S_RS14200 | WP_043898814.1 | iron ABC transporter substrate-binding protein |
| W5S_RS04290 | WP_012822705.1 | PTS sorbitol transporter subunit IIA |
| W5S_RS11185 | WP_014699967.1 | histidinol-phosphate aminotransferase 1 |
| W5S_RS15465 | WP_014700773.1 | phosphoribosylglycinamide formyltransferase |
| W5S_RS21455 | WP_033072536.1 | acetylornithine deacetylase |
| W5S_RS17380 | WP_014701120.1 | S-methyl-5-thioribose kinase |
| W5S_RS03960 | WP_014698779.1 | acetyl-coenzyme A synthetase |
| W5S_RS19715 | WP_014701559.1 | tRNA (adenosine(37)-N6)-dimethylallyltransferase MiaA |

| | | |
|---|---|---|
| W5S_RS18905 | WP_014701408.1 | nicotinate-nucleotide pyrophosphorylase |
| W5S_RS10690 | WP_014699878.1 | D-amino acid dehydrogenase small subunit |
| W5S_RS02035 | WP_014698573.1 | aminoimidazole riboside kinase |
| W5S_RS18020 | WP_014701230.1 | peptide-methionine (S)-S-oxide reductase |
| W5S_RS02205 | WP_025919470.1 | hypothetical protein |
| W5S_RS17875 | WP_005974672.1 | CTP synthetase |
| W5S_RS05495 | WP_014698916.1 | holo-ACP synthase |
| W5S_RS18990 | WP_043898838.1 | UDP-N-acetylmuramate--L-alanine ligase |
| W5S_RS19960 | WP_014701603.1 | homoserine O-succinyltransferase |
| W5S_RS01205 | WP_043898890.1 | bifunctional phosphoribosylaminoimidazolecarboxamide formyltransferase/inosine monophosphate cyclohydrolase |
| W5S_RS19470 | WP_014701513.1 | phosphoglycerate mutase gpmB |
| W5S_RS14865 | WP_005973959.1 | phosphoglyceromutase |
| W5S_RS18165 | WP_014701256.1 | pyrroline-5-carboxylate reductase |
| W5S_RS02800 | WP_014698671.1 | phosphoserine phosphatase |
| W5S_RS18620 | WP_014701350.1 | haloacid dehalogenase |
| W5S_RS04590 | WP_014698847.1 | cytosine permease |
| W5S_RS22370 | WP_014702026.1 | argininosuccinate synthase |
| W5S_RS01765 | WP_014698534.1 | argininosuccinate synthase |
| W5S_RS03915 | WP_014698773.1 | Ig family protein |
| W5S_RS20735 | WP_014701721.1 | alkaline phosphatase |
| W5S_RS01730 | WP_012822134.1 | hypothetical protein |
| W5S_RS01735 | WP_012822135.1 | aldolase |
| W5S_RS11530 | WP_014700040.1 | tagatose-bisphosphate aldolase |
| W5S_RS10655 | WP_014699872.1 | tagatose 1,6-diphosphate aldolase |
| W5S_RS01235 | WP_014698492.1 | amino acid ABC transporter permease |
| W5S_RS01240 | WP_014698493.1 | glutamate ABC transporter permease |
| W5S_RS15220 | WP_005976312.1 | glutamate/aspartate ABC transporter substrate-binding protein |
| W5S_RS23200 | WP_014702169.1 | amino acid ABC transporter permease |
| W5S_RS21040 | WP_014701782.1 | glutamate/glutamine/aspartate/asparagine ABC transporter substrate-binding protein |
| W5S_RS15230 | WP_014700733.1 | glutamate/aspartate ABC transporter permease GltK |
| W5S_RS15225 | WP_014700732.1 | glutamate/aspartate ABC transporter permease GltJ |
| W5S_RS01245 | WP_014698494.1 | glutamate/glutamine/aspartate/asparagine ABC transporter substrate-binding protein |
| W5S_RS21050 | WP_014701784.1 | polar amino acid ABC transporter permease |
| W5S_RS15235 | WP_005976306.1 | arginine ABC transporter ATP-binding protein |

| | | |
|---|---|---|
| W5S_RS13730 | WP_014700454.1 | acylphosphatase |
| W5S_RS03330 | WP_014698731.1 | transketolase |
| W5S_RS19620 | WP_014701541.1 | transketolase |
| W5S_RS04275 | WP_014698814.1 | transketolase |
| W5S_RS15190 | WP_014700726.1 | 2-octaprenyl-3-methyl-6-methoxy-1,4-benzoquinol hydroxylase |
| W5S_RS15135 | WP_014700721.1 | glucosamine-6-phosphate deaminase |
| W5S_RS03185 | WP_014698707.1 | altronate oxidoreductase |
| W5S_RS00320 | WP_014698400.1 | altronate oxidoreductase |
| W5S_RS16260 | WP_014700904.1 | UDP-3-O-(3-hydroxymyristoyl)glucosamine N-acyltransferase |
| W5S_RS16125 | WP_014700879.1 | mannonate dehydratase |
| W5S_RS20800 | WP_014701732.1 | MFS transporter |
| W5S_RS00335 | WP_014698402.1 | MFS transporter |
| W5S_RS00130 | WP_005976725.1 | MFS transporter |
| W5S_RS01695 | WP_012822128.1 | ADP-ribose diphosphatase |
| W5S_RS19910 | WP_014701593.1 | 2-oxoglutarate/malate translocator |
| W5S_RS14560 | WP_014700616.1 | phosphogluconate dehydrogenase (NADP(+)-dependent, decarboxylating) |
| W5S_RS00860 | WP_014698469.1 | agmatine deiminase |
| W5S_RS10525 | WP_014699847.1 | PTS mannose/fructose/sorbose transporter subunit IIC |
| W5S_RS10530 | WP_014699848.1 | PTS mannose transporter subunit EIIAB |
| W5S_RS10520 | WP_014699846.1 | PTS mannose transporter subunit IID |
| W5S_RS17715 | WP_014701176.1 | 2-C-methyl-D-erythritol 4-phosphate cytidylyltransferase |
| W5S_RS22950 | WP_005976539.1 | F0F1 ATP synthase subunit I |
| W5S_RS22990 | WP_005976558.1 | ATP synthase epsilon chain |
| W5S_RS22970 | WP_014702127.1 | F0F1 ATP synthase subunit delta |
| W5S_RS22975 | WP_005976551.1 | ATP synthase subunit alpha |
| W5S_RS22980 | WP_005976554.1 | ATP synthase subunit gamma |
| W5S_RS22985 | WP_014702128.1 | F0F1 ATP synthase subunit beta |
| W5S_RS20680 | WP_014701710.1 | Propanol dehydrogenase |
| W5S_RS14310 | WP_014700566.1 | adenosylmethionine--8-amino-7-oxononanoate aminotransferase BioA |
| W5S_RS14745 | WP_014700657.1 | 6-phosphogluconolactonase |
| W5S_RS01200 | WP_014698490.1 | phosphoribosylamine--glycine ligase |
| W5S_RS14120 | WP_014700528.1 | adenylate cyclase |
| W5S_RS06260 | WP_014699057.1 | acetylornithine deacetylase |
| W5S_RS17965 | WP_014701221.1 | CYTH domain-containing protein |
| W5S_RS20995 | WP_014701773.1 | adenylate cyclase |
| W5S_RS20415 | WP_014701660.1 | siroheme synthase |

| | | |
|---|---|---|
| W5S_RS17760 | WP_014701181.1 | siroheme synthase |
| W5S_RS07035 | WP_043898767.1 | uroporphyrinogen III methyltransferase |
| W5S_RS06635 | WP_014699124.1 | erythronate-4-phosphate dehydrogenase |
| W5S_RS12845 | WP_014700294.1 | riboflavin synthase |
| W5S_RS09605 | WP_014699684.1 | citrate (pro-3S)-lyase subunit beta |
| W5S_RS09610 | WP_014699685.1 | citrate lyase subunit alpha |
| W5S_RS09600 | WP_014699683.1 | citrate lyase ACP |
| W5S_RS05150 | WP_012822848.1 | ribonucleotide-diphosphate reductase subunit beta |
| W5S_RS15590 | WP_005976154.1 | ribonucleoside-diphosphate reductase subunit alpha |
| W5S_RS15595 | WP_014700796.1 | ribonucleotide-diphosphate reductase subunit beta |
| W5S_RS05160 | WP_005968995.1 | class Ib ribonucleoside-diphosphate reductase assembly flavoprotein NrdI |
| W5S_RS05155 | WP_014698893.1 | ribonucleotide-diphosphate reductase |
| W5S_RS12880 | WP_014700300.1 | aspartate racemase |
| W5S_RS18330 | WP_014701290.1 | alanine racemase |
| W5S_RS08385 | WP_014699463.1 | aspartate racemase |
| W5S_RS15325 | WP_014700748.1 | aspartate racemase |
| W5S_RS08260 | WP_014699440.1 | aspartate racemase |
| W5S_RS22630 | WP_014702077.1 | glutamate dehydrogenase |
| W5S_RS00890 | WP_012822006.1 | glycerol kinase |
| W5S_RS12815 | WP_010276601.1 | lactoylglutathione lyase |
| W5S_RS02155 | WP_010298253.1 | aspartate carbamoyltransferase regulatory subunit |
| W5S_RS12860 | WP_014700296.1 | L-lactate permease |
| W5S_RS16825 | WP_014701009.1 | arsenical efflux pump membrane protein ArsB |
| W5S_RS05610 | WP_014698936.1 | nitric oxide dioxygenase |
| W5S_RS18965 | WP_043899162.1 | UDP-3-O-[3-hydroxymyristoyl N-acetylglucosamine deacetylase |
| W5S_RS14695 | WP_014700647.1 | dCTP deaminase |
| W5S_RS19345 | WP_014701493.1 | carbamoyl-phosphate synthase small subunit |
| W5S_RS19340 | WP_014701492.1 | carbamoyl phosphate synthase large subunit |
| W5S_RS13185 | WP_025918561.1 | ammonia-forming cytochrome c nitrite reductase subunit c552 |
| W5S_RS14340 | WP_014700573.1 | NAD-dependent malic enzyme |
| W5S_RS17885 | WP_014701204.1 | GTP diphosphokinase |
| W5S_RS04485 | WP_012822741.1 | fumarate hydratase |
| W5S_RS11130 | WP_014699957.1 | class II fumarate hydratase |
| W5S_RS09690 | WP_014699698.1 | 3-deoxy-manno-octulosonate cytidylyltransferase |
| W5S_RS19160 | WP_014701457.1 | acetolactate synthase isozyme 1 small subunit |
| W5S_RS19055 | WP_014701436.1 | acetolactate synthase small subunit |
| W5S_RS21270 | WP_005969103.1 | acetolactate synthase 2 small subunit |
| W5S_RS19155 | WP_014701456.1 | acetolactate synthase |

| | | |
|---|---|---|
| W5S_RS19060 | WP_014701437.1 | acetolactate synthase 3 large subunit |
| W5S_RS21275 | WP_014701819.1 | acetolactate synthase 2 catalytic subunit |
| W5S_RS03640 | WP_012822565.1 | acetolactate synthase AlsS |
| W5S_RS19435 | WP_014701508.1 | threonine synthase |
| W5S_RS17430 | WP_014701130.1 | aminotransferase |
| W5S_RS14835 | WP_014700673.1 | galactose-1-epimerase |
| W5S_RS09125 | WP_014699598.1 | Lysine-specific permease |
| W5S_RS17520 | WP_014701145.1 | glycine betaine/L-proline ABC transporter ATP-binding protein |
| W5S_RS23010 | WP_014702132.1 | glycine/betaine ABC transporter ATP-binding protein |
| W5S_RS17530 | WP_025920047.1 | glycine betaine ABC transporter substrate-binding protein |
| W5S_RS23025 | WP_014702135.1 | glycine/betaine ABC transporter substrate-binding protein |
| W5S_RS03290 | WP_014698724.1 | kojibiose phosphorylase |
| W5S_RS02190 | WP_012822210.1 | N-acetyltransferase |
| W5S_RS00855 | WP_012821999.1 | N-carbamoylputrescine amidase |
| W5S_RS03190 | WP_014698708.1 | altronate hydrolase |
| W5S_RS09230 | WP_014699618.1 | 2-deoxyribose-5-phosphate aldolase |
| W5S_RS17925 | WP_014701213.1 | MFS transporter |
| W5S_RS01035 | WP_012822025.1 | UDP-N-acetylenolpyruvoylglucosamine reductase |
| W5S_RS04335 | WP_012822714.1 | coproporphyrinogen III oxidase |
| W5S_RS06880 | WP_014699160.1 | coproporphyrinogen III oxidase |
| W5S_RS14890 | WP_014700683.1 | zinc transporter ZitB |
| W5S_RS10335 | WP_014699816.1 | aromatic amino acid transporter |
| W5S_RS04325 | WP_014698820.1 | tryptophan permease |
| W5S_RS18885 | WP_014701404.1 | aromatic amino acid transporter AroP |
| W5S_RS08255 | WP_014699439.1 | pectin lyase |
| W5S_RS10815 | WP_005968735.1 | formyltetrahydrofolate deformylase |
| W5S_RS15545 | WP_043898820.1 | isochorismate synthase MenF |
| W5S_RS09470 | WP_014699658.1 | formate transporter FocA |
| W5S_RS11180 | WP_014699966.1 | NAD(P)-dependent oxidoreductase |
| W5S_RS01145 | WP_014698484.1 | sulfur carrier protein ThiS |
| W5S_RS01140 | WP_012822036.1 | thiazole synthase |
| W5S_RS01150 | WP_014698485.1 | adenylyltransferase ThiF |
| W5S_RS07050 | WP_014699194.1 | acireductone dioxygenase |
| W5S_RS17415 | WP_014701127.1 | acireductone dioxygenase |
| W5S_RS16800 | WP_014701003.1 | phosphopyruvate hydratase |
| W5S_RS17870 | WP_005974669.1 | enolase |
| W5S_RS14465 | WP_014700597.1 | 5-dehydro-2-deoxygluconokinase |
| W5S_RS11620 | WP_071531107.1 | ribose-phosphate pyrophosphokinase |

| | | |
|---|---|---|
| W5S_RS04440 | WP_005971895.1 | sulfate transporter CysZ |
| W5S_RS05335 | WP_012822880.1 | ABC transporter permease |
| W5S_RS05345 | WP_012822881.1 | sulfate ABC transporter permease subunit CysW |
| W5S_RS05340 | WP_005969061.1 | sulfate ABC transporter permease subunit CysT |
| W5S_RS05350 | WP_005969065.1 | sulfate ABC transporter ATP-binding protein |
| W5S_RS04380 | WP_014698826.1 | sulfate ABC transporter substrate-binding protein |
| W5S_RS04390 | WP_014698828.1 | Sulfate ABC transporter inner membrane subunit CysW |
| W5S_RS04385 | WP_014698827.1 | sulfate ABC transporter permease CysT |
| W5S_RS19410 | WP_014701504.1 | MFS transporter |
| W5S_RS19390 | WP_014701501.1 | Na+/H+ antiporter NhaA |
| W5S_RS14335 | WP_014700572.1 | cytidine deaminase |
| W5S_RS21865 | WP_014701929.1 | uridine phosphorylase |
| W5S_RS13085 | WP_014700338.1 | periplasmic nitrate reductase subunit alpha |
| W5S_RS12320 | WP_014700196.1 | respiratory nitrate reductase subunit gamma |
| W5S_RS12325 | WP_014700197.1 | nitrate reductase molybdenum cofactor assembly chaperone |
| W5S_RS12330 | WP_014700198.1 | nitrate reductase subunit beta |
| W5S_RS12335 | WP_014700199.1 | nitrate reductase subunit alpha |
| W5S_RS04205 | WP_014698806.1 | manganese transporter MntH |
| W5S_RS12480 | WP_014700229.1 | zinc transporter ZntB |
| W5S_RS20945 | WP_014701761.1 | magnesium transporter CorA |
| W5S_RS13105 | WP_005970951.1 | cytochrome c-type protein NapC |
| W5S_RS13100 | WP_005970953.1 | nitrate reductase cytochrome C550 subunit |
| W5S_RS13090 | WP_014700339.1 | ferredoxin-type protein NapG |
| W5S_RS13095 | WP_014700340.1 | quinol dehydrogenase ferredoxin subunit NapH |
| W5S_RS03700 | WP_014698754.1 | glycoside hydrolase |
| W5S_RS17620 | WP_014701164.1 | MFS transporter |
| W5S_RS00560 | WP_014698427.1 | 3-hydroxyisobutyrate dehydrogenase |
| W5S_RS04320 | WP_012822711.1 | NADP-dependent malic enzyme |
| W5S_RS12635 | WP_014700257.1 | oxidoreductase subunit alpha |
| W5S_RS22420 | WP_014702037.1 | sulfurtransferase FdhD |
| W5S_RS13470 | WP_014700406.1 | thiamine kinase |
| W5S_RS09485 | WP_014699661.1 | 3-phosphoshikimate 1-carboxyvinyltransferase |
| W5S_RS03820 | WP_043898746.1 | lipid A biosynthesis palmitoleoyl acyltransferase |
| W5S_RS13625 | WP_014700432.1 | lipid A biosynthesis lauroyl acyltransferase |
| W5S_RS20345 | WP_025919614.1 | glutamine amidotransferase |
| W5S_RS20450 | WP_014701666.1 | phosphoglycolate phosphatase |
| W5S_RS12185 | WP_014700170.1 | phosphoglycolate phosphatase |
| W5S_RS20865 | WP_014701744.1 | glycosyl hydrolase family 3 |

| | | |
|---|---|---|
| W5S_RS01355 | WP_012822068.1 | 3-dehydroquinate dehydratase |
| W5S_RS22570 | WP_014702065.1 | glutathione-disulfide reductase |
| W5S_RS13490 | WP_014700410.1 | PTS glucose transporter subunit IIBC |
| W5S_RS04420 | WP_005971886.1 | PTS glucose transporter subunit IIA |
| W5S_RS15540 | WP_014700787.1 | 2-succinyl-5-enolpyruvyl-6-hydroxy-3-cyclohexene-1-carboxylic-acid synthase |
| W5S_RS06685 | WP_014699130.1 | 3-octaprenyl-4-hydroxybenzoate carboxy-lyase |
| W5S_RS21385 | WP_014701835.1 | 3-octaprenyl-4-hydroxybenzoate decarboxylase |
| W5S_RS02915 | WP_012822336.1 | phosphonopyruvate decarboxylase |
| W5S_RS01010 | WP_012822024.1 | glutamate racemase |
| W5S_RS10735 | WP_014699887.1 | peptide-methionine (R)-S-oxide reductase |
| W5S_RS04030 | WP_014698785.1 | D-mannonate oxidoreductase |
| W5S_RS04565 | WP_014698845.1 | D-mannonate oxidoreductase |
| W5S_RS06245 | WP_014699055.1 | cytochrome c552 |
| W5S_RS06250 | WP_014699056.1 | D-alanine--poly(phosphoribitol) ligase |
| W5S_RS12235 | WP_014700179.1 | NAD-dependent succinate-semialdehyde dehydrogenase |
| W5S_RS19905 | WP_014701592.1 | Inorganic phosphate transporter sodium-dependent |
| W5S_RS14885 | WP_014700682.1 | 3-deoxy-7-phosphoheptulonate synthase |
| W5S_RS13310 | WP_014700376.1 | 3-deoxy-7-phosphoheptulonate synthase |
| W5S_RS05045 | WP_012822836.1 | phospho-2-dehydro-3-deoxyheptonate aldolase |
| W5S_RS12715 | WP_014700272.1 | transketolase |
| W5S_RS12720 | WP_014700273.1 | transketolase |
| W5S_RS19920 | WP_014701595.1 | methionine synthase |
| W5S_RS15920 | WP_014700844.1 | 1-deoxy-D-xylulose-5-phosphate synthase |
| W5S_RS19505 | WP_014701519.1 | D-3-phosphoglycerate dehydrogenase |
| W5S_RS13015 | WP_014700325.1 | D-isomer specific 2-hydroxyacid dehydrogenase |
| W5S_RS06060 | WP_014699019.1 | galactose-1-phosphate uridylyltransferase |
| W5S_RS05575 | WP_014698930.1 | phosphoribosylformylglycinamidine synthase |
| W5S_RS14460 | WP_014700596.1 | myo-inosose-2 dehydratase |
| W5S_RS03650 | WP_012822567.1 | glycine cleavage system protein T |
| W5S_RS22020 | WP_014701961.1 | 3-deoxy-D-manno-octulosonic acid transferase |
| W5S_RS06860 | WP_014699156.1 | PTS trehalose transporter subunit IIBC |
| W5S_RS05210 | WP_012822858.1 | aspartate 1-decarboxylase |
| W5S_RS22875 | WP_005976509.1 | D-ribose ABC transporter substrate-binding protein |
| W5S_RS12725 | WP_014700274.1 | sugar ABC transporter permease |
| W5S_RS19550 | WP_014701528.1 | ABC transporter permease |
| W5S_RS22885 | WP_014702114.1 | ribose ABC transporter ATP-binding protein RbsA |
| W5S_RS22880 | WP_005976511.1 | ribose ABC transporter permease |
| W5S_RS22890 | WP_014702115.1 | D-ribose pyranase |

| | | |
|---|---|---|
| W5S_RS12740 | WP_005972510.1 | D-ribose ABC transporter substrate-binding protein |
| W5S_RS09140 | WP_014699601.1 | sugar ABC transporter permease |
| W5S_RS09145 | WP_014699602.1 | ribose import ATP-binding protein RbsA |
| W5S_RS12730 | WP_043899068.1 | sugar ABC transporter ATP-binding protein |
| W5S_RS08440 | WP_014699477.1 | 1-pyrroline dehydrogenase |
| W5S_RS16415 | WP_014700931.1 | biotin sulfoxide reductase |
| W5S_RS23215 | WP_014702172.1 | Xanthine/uracil/thiamine/ascorbate permease |
| W5S_RS15895 | WP_014700840.1 | 2-dehydropantoate 2-reductase |
| W5S_RS16875 | WP_014701020.1 | sodium-independent anion transporter |
| W5S_RS07080 | WP_014699201.1 | Antisigma-factor antagonist, STAS |
| W5S_RS03465 | WP_012822537.1 | phosphoglucosamine mutase |
| W5S_RS21945 | WP_014701943.1 | L-threonine 3-dehydrogenase |
| W5S_RS11155 | WP_014699962.1 | dethiobiotin synthase |
| W5S_RS14290 | WP_014700562.1 | dethiobiotin synthase |
| W5S_RS02920 | WP_014698688.1 | 2-hydroxy-3-oxopropionate reductase |
| W5S_RS14580 | WP_014700620.1 | NAD(P)-dependent oxidoreductase |
| W5S_RS18865 | WP_014701401.1 | dihydrolipoyl dehydrogenase |
| W5S_RS15925 | WP_014700845.1 | thiamine-monophosphate kinase |
| W5S_RS13350 | WP_014700383.1 | D-gluconate kinase |
| W5S_RS14495 | WP_014700603.1 | 3D-(3,5/4)-trihydroxycyclohexane-1,2-dione acylhydrolase (decyclizing) |
| W5S_RS13855 | WP_014700478.1 | MFS transporter |
| W5S_RS15510 | WP_015730946.1 | tyrosine transporter TyrP |
| W5S_RS12385 | WP_014700211.1 | dihydromonapterin reductase |
| W5S_RS20925 | WP_014701757.1 | phospholipase A |
| W5S_RS02575 | WP_012822274.1 | L-rhamnose isomerase |
| W5S_RS21420 | WP_014701841.1 | ubiquinone biosynthesis regulatory protein kinase UbiB |
| W5S_RS21350 | WP_005973457.1 | potassium transporter |
| W5S_RS20045 | WP_014701615.1 | Trk system potassium transport protein TrkA |
| W5S_RS10045 | WP_005967769.1 | glucose-6-phosphate dehydrogenase |
| W5S_RS17300 | WP_014701107.1 | gamma-glutamyl-phosphate reductase |
| W5S_RS23230 | WP_014702174.1 | oxidoreductase FeS-binding subunit |
| W5S_RS01595 | WP_012822109.1 | glutamate synthase subunit beta |
| W5S_RS01600 | WP_014698517.1 | glutamate synthase large subunit |
| W5S_RS10490 | WP_014699841.1 | hypothetical protein |
| W5S_RS10485 | WP_014699840.1 | membrane protein |
| W5S_RS10495 | WP_005968833.1 | metal ABC transporter substrate-binding protein |
| W5S_RS10480 | WP_014699839.1 | iron ABC transporter permease |
| W5S_RS17420 | WP_014701128.1 | acireductone synthase |

| | | |
|---|---|---|
| W5S_RS11305 | WP_014699994.1 | transport secretion system IV protein, VirB6 |
| W5S_RS19005 | WP_014701429.1 | UDP-N-acetylmuramoyl-L-alanine--D-glutamate ligase |
| W5S_RS20820 | WP_014701736.1 | glycogen phosphorylase |
| W5S_RS11110 | WP_014699953.1 | adenosine deaminase |
| W5S_RS20505 | WP_014701678.1 | carboxypeptidase/penicillin-binding protein 1A |
| W5S_RS05260 | WP_012822867.1 | bifunctional glycosyl transferase/transpeptidase |
| W5S_RS00120 | WP_012821883.1 | 2-dehydro-3-deoxy-6-phosphogalactonate aldolase |
| W5S_RS15520 | WP_014700784.1 | 2-succinylbenzoate-CoA ligase |
| W5S_RS02860 | WP_014698678.1 | isochorismate synthase EntC |
| W5S_RS17980 | WP_014701224.1 | undecaprenyl-diphosphatase |
| W5S_RS15450 | WP_014700771.1 | uracil/xanthine transporter |
| W5S_RS21430 | WP_005973427.1 | bifunctional demethylmenaquinone methyltransferase/2-methoxy-6-polyprenyl-1,4-benzoquinol methylase |
| W5S_RS00910 | WP_012822008.1 | 1,4-dihydroxy-2-naphthoate polyprenyltransferase |
| W5S_RS02410 | WP_014698628.1 | ABC transporter substrate-binding protein |
| W5S_RS16080 | WP_014700869.1 | L-asparaginase |
| W5S_RS20590 | WP_014701695.1 | citrate (Si)-synthase |
| W5S_RS15020 | WP_005973887.1 | citrate (Si)-synthase |
| W5S_RS10395 | WP_014699824.1 | NAD(+) synthetase |
| W5S_RS16250 | WP_014700903.1 | acyl-[acyl-carrier-protein--UDP-N-acetylglucosamine O-acyltransferase |
| W5S_RS03075 | WP_012822479.1 | aspartate ammonia-lyase |
| W5S_RS02570 | WP_012822273.1 | rhamnulose-1-phosphate aldolase |
| W5S_RS06335 | WP_014699068.1 | mechanosensitive ion channel protein MscS |
| W5S_RS19835 | WP_014701579.1 | miniconductance mechanosensitive channel MscM |
| W5S_RS21190 | WP_014701805.1 | thiol reductase thioredoxin |
| W5S_RS08205 | WP_014699429.1 | MFS transporter |
| W5S_RS03695 | WP_014698753.1 | MFS transporter |
| W5S_RS12665 | WP_014700263.1 | MFS transporter |
| W5S_RS21450 | WP_014701846.1 | N-acetyl-gamma-glutamyl-phosphate reductase |
| W5S_RS06140 | WP_014699035.1 | UDP-2,3-diacylglucosamine diphosphatase |
| W5S_RS05215 | WP_012822859.1 | pantoate--beta-alanine ligase |
| W5S_RS17800 | WP_014701189.1 | sulfite reductase subunit alpha |
| W5S_RS17795 | WP_014701188.1 | sulfite reductase subunit beta |
| W5S_RS03060 | WP_014698700.1 | protein-disulfide reductase DsbD |
| W5S_RS12020 | WP_014700138.1 | allophanate hydrolase |
| W5S_RS15040 | WP_014700704.1 | allophanate hydrolase |
| W5S_RS15045 | WP_014700705.1 | Kinase autophosphorylation inhibitor kipI |
| W5S_RS00880 | WP_012822004.1 | ferredoxin--NADP(+) reductase |

| | | |
|---|---|---|
| W5S_RS21955 | WP_014701945.1 | ADP-L-glycero-D-mannoheptose-6-epimerase |
| W5S_RS22060 | WP_005967975.1 | deoxyuridine 5'-triphosphate nucleotidohydrolase |
| W5S_RS02635 | WP_012822285.1 | rhamnose/proton symporter RhaT |
| W5S_RS15670 | WP_014700807.1 | ferrochelatase |
| W5S_RS13505 | WP_014700413.1 | dTMP kinase |
| W5S_RS19145 | WP_014701454.1 | thiamine ABC transporter ATP-binding protein |
| W5S_RS19135 | WP_014701452.1 | thiamine transporter substrate binding subunit |
| W5S_RS19140 | WP_014701453.1 | thiamine/thiamine pyrophosphate ABC transporter permease ThiP |
| W5S_RS05125 | WP_014698890.1 | hydroxyacylglutathione hydrolase |
| W5S_RS10055 | WP_014699765.1 | phosphoribosylglycinamide formyltransferase 2 |
| W5S_RS21950 | WP_014701944.1 | glycine C-acetyltransferase |
| W5S_RS02180 | WP_012822209.1 | ornithine carbamoyltransferase |
| W5S_RS00045 | WP_014698374.1 | PTS cellobiose IIC component |
| W5S_RS00160 | WP_012821891.1 | PTS sugar transporter subunit IIB |
| W5S_RS10000 | WP_014699756.1 | PTS cellobiose IIC component |
| W5S_RS09995 | WP_043898785.1 | molecular chaperone TorD |
| W5S_RS17770 | WP_014701183.1 | PTS sugar transporter subunit IIB |
| W5S_RS17775 | WP_014701184.1 | PTS cellobiose IIC component |
| W5S_RS00165 | WP_012821892.1 | PTS cellobiose transporter subunit IIC |
| W5S_RS00050 | WP_012821870.1 | molecular chaperone TorD |
| W5S_RS00040 | WP_012821868.1 | PTS sugar transporter subunit IIB |
| W5S_RS05280 | WP_012822871.1 | membrane protein |
| W5S_RS03050 | WP_012822474.1 | lysine transporter LysE |
| W5S_RS20315 | WP_014701640.1 | alcohol dehydrogenase |
| W5S_RS09480 | WP_014699660.1 | phosphoserine transaminase |
| W5S_RS20640 | WP_014701704.1 | aldehyde dehydrogenase |
| W5S_RS12885 | WP_014700301.1 | 4-oxalocrotonate tautomerase family enzyme |
| W5S_RS11190 | WP_014699968.1 | 4-oxalocrotonate tautomerase |
| W5S_RS04220 | WP_012822691.1 | tautomerase |
| W5S_RS20620 | WP_005969393.1 | glycyl-radical enzyme activating protein |
| W5S_RS17440 | WP_014701132.1 | glycyl-radical enzyme activating protein family |
| W5S_RS18695 | WP_014701365.1 | choline TMA-lyase-activating enzyme |
| W5S_RS09460 | WP_005967552.1 | pyruvate formate lyase 1-activating protein |
| W5S_RS09405 | WP_014699647.1 | thioredoxin-disulfide reductase |
| W5S_RS15335 | WP_014700750.1 | succinyl-diaminopimelate desuccinylase |
| W5S_RS12015 | WP_014700137.1 | urea carboxylase |
| W5S_RS12915 | WP_014700307.1 | urea carboxylase |
| W5S_RS09750 | WP_014699710.1 | porin |
| W5S_RS10225 | WP_005968924.1 | NUDIX hydrolase |

| | | |
|---|---|---|
| W5S_RS13795 | WP_014700467.1 | beta-hydroxydecanoyl-ACP dehydratase |
| W5S_RS16255 | WP_005975923.1 | beta-hydroxyacyl-ACP dehydratase |
| W5S_RS19580 | WP_014701534.1 | carbohydrate kinase |
| W5S_RS13530 | WP_005970506.1 | acyl carrier protein |
| W5S_RS18210 | WP_014701264.1 | acyl-[ACP--phospholipid O-acyltransferase |
| W5S_RS21140 | WP_014701796.1 | 4-alpha-L-fucosyltransferase |
| W5S_RS13405 | WP_014700395.1 | N-acetylglucosamine kinase |
| W5S_RS05555 | WP_014698926.1 | N-acetylmuramic acid 6-phosphate etherase |
| W5S_RS21130 | WP_014701794.1 | lipopolysaccharide N-acetylmannosaminouronosyltransferase |
| W5S_RS12050 | WP_043898799.1 | porin |
| W5S_RS00315 | WP_014698399.1 | amino-acid metabolite efflux pump |
| W5S_RS03950 | WP_012822639.1 | cation acetate symporter |
| W5S_RS18900 | WP_014701407.1 | nucleoside-specific channel-forming protein Tsx |
| W5S_RS18895 | WP_014701406.1 | N-acetylmuramoyl-L-alanine amidase |
| W5S_RS13465 | WP_014700405.1 | beta-hexosaminidase |
| W5S_RS15855 | WP_014700835.1 | AmpG family muropeptide MFS transporter |
| W5S_RS14855 | WP_014700677.1 | hypothetical protein |
| W5S_RS14860 | WP_014700678.1 | CusA/CzcA family heavy metal efflux RND transporter |
| W5S_RS07090 | WP_014699203.1 | RND efflux system, hypothetical protein, NodT family |
| W5S_RS14845 | WP_014700675.1 | cation transporter |
| W5S_RS06350 | WP_014699071.1 | alanine transaminase |
| W5S_RS18715 | WP_014701369.1 | allose kinase |
| W5S_RS17460 | WP_014701136.1 | Zn-dependent hydrolase |
| W5S_RS03985 | WP_014698783.1 | amidohydrolase/deacetylase family metallohydrolase |
| W5S_RS01790 | WP_012822144.1 | 2,5-didehydrogluconate reductase A |
| W5S_RS10720 | WP_014699884.1 | aldo/keto reductase |
| W5S_RS00675 | WP_012821980.1 | sugar transporter |
| W5S_RS19535 | WP_014701524.1 | Arginine exporter protein ArgO |
| W5S_RS16600 | WP_014700964.1 | amino acid permease |
| W5S_RS01510 | WP_012822095.1 | calcium/sodium antiporter |
| W5S_RS00680 | WP_012821981.1 | cation-efflux pump FieF |
| W5S_RS08380 | WP_014699462.1 | D-cysteine desulfhydrase |
| W5S_RS16400 | WP_014700928.1 | cysteine sulfinate desulfinase |
| W5S_RS02870 | WP_043898734.1 | 2,3-dihydro-2,3-dihydroxybenzoate dehydrogenase |
| W5S_RS02865 | WP_014698679.1 | (2,3-dihydroxybenzoyl)adenylate synthase |
| W5S_RS18945 | WP_014701418.1 | 8-oxo-dGTP diphosphatase |
| W5S_RS09965 | WP_014699749.1 | NUDIX pyrophosphatase |
| W5S_RS21145 | WP_014701797.1 | lipid III flippase WzxE |

| | | |
|---|---|---|
| W5S_RS02845 | WP_012822320.1 | 4'-phosphopantetheinyl transferase |
| W5S_RS18790 | WP_014701385.1 | enterochelin esterase |
| W5S_RS15760 | WP_014700820.1 | acyl-CoA thioesterase II |
| W5S_RS21380 | WP_014701834.1 | NAD(P)H-flavin reductase |
| W5S_RS00325 | WP_014698401.1 | Starvation-sensing protein RspB |
| W5S_RS21310 | WP_014701827.1 | ABC transporter substrate-binding protein |
| W5S_RS21295 | WP_014701824.1 | sugar ABC transporter permease YjfF |
| W5S_RS21305 | WP_014701826.1 | sugar ABC transporter ATP-binding protein |
| W5S_RS21300 | WP_014701825.1 | ABC transporter permease |
| W5S_RS04225 | WP_014698808.1 | GDP-mannose pyrophosphatase NudK |
| W5S_RS22625 | WP_014702076.1 | Aminophosphonate oxidoreductase |
| W5S_RS20835 | WP_014701739.1 | glycogen debranching enzyme |
| W5S_RS15125 | WP_014700719.1 | glutamine--tRNA ligase |
| W5S_RS13865 | WP_014700480.1 | threonine/homoserine exporter RhtA |
| W5S_RS20910 | WP_014701754.1 | homoserine/homoserine lactone efflux protein |
| W5S_RS19585 | WP_014701535.1 | 3-keto-L-gulonate-6-phosphate decarboxylase ulaD |
| W5S_RS06200 | WP_014699046.1 | 4-amino-4-deoxy-L-arabinose lipid A transferase |
| W5S_RS12490 | WP_014700231.1 | murein peptide amidase A |
| W5S_RS21910 | WP_014701937.1 | phospholipid:lipid A palmitoyltransferase |
| W5S_RS15275 | WP_014700741.1 | penicillin-binding protein 2 |
| W5S_RS19025 | WP_014701432.1 | peptidoglycan glycosyltransferase FtsI |
| W5S_RS15290 | WP_014700744.1 | serine-type D-Ala-D-Ala carboxypeptidase |
| W5S_RS09220 | WP_014699616.1 | D-alanyl-D-alanine carboxypeptidase |
| W5S_RS03435 | WP_012822534.1 | serine-type D-Ala-D-Ala carboxypeptidase |
| W5S_RS14110 | WP_014700526.1 | D-alanyl-D-alanine endopeptidase |
| W5S_RS06590 | WP_014699116.1 | penicillin-insensitive murein endopeptidase |
| W5S_RS16170 | WP_014700889.1 | murein transglycosylase B |
| W5S_RS16560 | WP_033071863.1 | lytic murein transglycosylase |
| W5S_RS19485 | WP_014701515.1 | murein transglycosylase |
| W5S_RS16420 | WP_014700932.1 | murein transglycosylase A |
| W5S_RS05570 | WP_043898758.1 | lytic transglycosylase F |
| W5S_RS14135 | WP_014700531.1 | S-methylmethionine permease |
| W5S_RS01170 | WP_014698488.1 | NADH pyrophosphatase |
| W5S_RS01725 | WP_014698531.1 | Modulator of drug activity B |
| W5S_RS04475 | WP_014698833.1 | anaerobic nitric oxide reductase flavorubredoxin |
| W5S_RS04470 | WP_012822738.1 | NADH:flavorubredoxin oxidoreductase |
| W5S_RS12340 | WP_014700201.1 | NarK family nitrate/nitrite MFS transporter |
| W5S_RS06770 | WP_014699145.1 | 5'-deoxynucleotidase |
| W5S_RS19825 | WP_014701577.1 | ribosome small subunit-dependent GTPase |

| | | |
|---|---|---|
| W5S_RS17880 | WP_014701203.1 | nucleoside triphosphate pyrophosphohydrolase |
| W5S_RS18180 | WP_014701259.1 | non-canonical purine NTP pyrophosphatase, RdgB/HAM1 family |
| W5S_RS14765 | WP_014700659.1 | pyridoxal phosphatase |
| W5S_RS21090 | WP_014701790.1 | glucose-1-phosphatase |
| W5S_RS06090 | WP_014699025.1 | exopolyphosphatase |
| W5S_RS17685 | WP_014701172.1 | proline--tRNA ligase |
| W5S_RS20900 | WP_014701752.1 | sugar/pyridoxal phosphate phosphatase YigL |
| W5S_RS03980 | WP_014698782.1 | SelA-like pyridoxal phosphate-dependent enzyme |
| W5S_RS03200 | WP_014698709.1 | serine/threonine transporter SstT |
| W5S_RS19680 | WP_014701553.1 | UDP-N-acetylmuramate:L-alanyl-gamma-D-glutamyl-meso-diaminopimelate ligase |
| W5S_RS06195 | WP_014699045.1 | 4-deoxy-4-formamido-L-arabinose-phosphoundecaprenol deformylase |
| W5S_RS21010 | WP_014701776.1 | uroporphyrinogen-III C-methyltransferase |
| W5S_RS09855 | WP_014699730.1 | glutaredoxin 2 |
| W5S_RS16820 | WP_014701008.1 | arsenate reductase (glutaredoxin) |
| W5S_RS22815 | WP_014702107.1 | protein disulfide oxidoreductase DsbA |
| W5S_RS10635 | WP_014699868.1 | disulfide bond formation protein B |
| W5S_RS03775 | WP_043898744.1 | bifunctional protein-disulfide isomerase/oxidoreductase DsbC |
| W5S_RS21135 | WP_014701795.1 | enterobacterial common antigen polymerase |
| W5S_RS21175 | WP_014701803.1 | polysaccharide chain length modulation protein |
| W5S_RS12825 | WP_014700290.1 | glutaredoxin |
| W5S_RS09245 | WP_005967344.1 | glutaredoxin, GrxA family |
| W5S_RS21970 | WP_014701949.1 | Lipid A-core:surface polymer ligase WaaL |
| W5S_RS06180 | WP_014699042.1 | UDP-4-amino-4-deoxy-L-arabinose--oxoglutarate aminotransferase |
| W5S_RS06185 | WP_014699043.1 | undecaprenyl-phosphate 4-deoxy-4-formamido-L-arabinose transferase |
| W5S_RS11500 | WP_014700034.1 | PTS N-acetylgalactosamine transporter subunit IIB |
| W5S_RS11505 | WP_014700035.1 | PTS N-acetylgalactosamine transporter subunit IIC |
| W5S_RS11515 | WP_014700037.1 | PTS N-acetylgalactosamine transporter subunit IIA |
| W5S_RS11520 | WP_014700038.1 | N-acetylglucosamine-6-phosphate deacetylase |
| W5S_RS04435 | pseudogene | cysteine synthase A |
| W5S_RS15260 | pseudogene | nicotinate-nicotinamide nucleotide adenylyltransferase |
| W5S_RS18240 | pseudogene | glycerate kinase |
| W5S_RS16640 | pseudogene | IS256 family transposase |
| W5S_RS03255 | pseudogene | carbohydrate porin |
| W5S_RS13210 | pseudogene | carbohydrate porin |

| | | |
|---|---|---|
| W5S_RS11870 | pseudogene | ABC transporter substrate-binding protein |
| W5S_RS09445 | pseudogene | cytosine permease |
| W5S_RS21960 | pseudogene | ADP-heptose--LPS heptosyltransferase |
| W5S_RS16735 | pseudogene | hypothetical protein |
| W5S_RS14585 | pseudogene | UDP-galactopyranose mutase |
| W5S_RS23020 | pseudogene | ABC transporter permease |
| W5S_RS10030 | pseudogene | lauroyl-Kdo(2)-lipid IV(A) myristoyltransferase |
| W5S_RS10550 | pseudogene | aminodeoxychorismate synthase component I |
| W5S_RS05805 | pseudogene | IMP dehydrogenase |
| W5S_RS03295 | pseudogene | PTS trehalose transporter subunit IIBC |
| W5S_RS19545 | pseudogene | sugar ABC transporter ATP-binding protein |
| W5S_RS07830 | pseudogene | hypothetical protein |
| W5S_RS02490 | pseudogene | aspartate ammonia-lyase |
| W5S_RS11510 | pseudogene | PTS N-acetylgalactosamine transporter subunit IID |
| W5S_RS12800 | pseudogene | anhydro-N-acetylmuramic acid kinase |

**Supplementary Table 3:** Biomass Composition

| COMPOUND NAME | ID | AMOUNT |
|---|---|---|
| Co2+ | cpd00149 | 0 |
| L-Asparagine | cpd00132 | 0,2 |
| DNA replication | cpd17042 | 1 |
| Menaquinone 8 | cpd15500 | 0 |
| Pyridoxal phosphate | cpd00016 | 0 |
| CTP | cpd00052 | 0,08 |
| H2O | cpd00001 | 35,54 |
| Protein biosynthesis | cpd17041 | 1 |
| Putrescine | cpd00118 | 0 |
| L-Arginine | cpd00051 | 0,25 |
| Sulfate | cpd00048 | 0 |
| ATP | cpd00002 | 40,11 |
| 2 Demethylmenaquinone 8 | cpd15352 | 0 |
| dCTP | cpd00356 | 0,02 |
| TTP | cpd00357 | 0,02 |
| Dianteisoheptadecanoylphosphatidylethanolamine | cpd15696 | 0,01 |
| Anteisoheptadecanoylcardiolipin B subtilis | cpd15795 | 0,01 |
| Diisoheptadecanoylphosphatidylethanolamine | cpd15695 | 0,01 |
| Diisoheptadecanoylphosphatidylglycerol | cpd15722 | 0,01 |
| L-Aspartate | cpd00041 | 0,2 |
| S-Adenosyl-L-methionine | cpd00017 | 0 |
| L-Isoleucine | cpd00322 | 0,24 |
| L-Methionine | cpd00060 | 0,13 |
| dGTP | cpd00241 | 0,02 |
| 5-Methyltetrahydrofolate | cpd00345 | 0 |
| GSH | cpd00042 | 0 |
| Cu2+ | cpd00058 | 0 |
| UTP | cpd00062 | 0,09 |
| Fe3+ | cpd10516 | 0 |
| Phosphatidylglycerol dioctadecanoyl | cpd15540 | 0,01 |
| Mn2+ | cpd00030 | 0 |
| Tetrahydrofolate | cpd00087 | 0 |
| TPP | cpd00056 | 0 |
| K+ | cpd00205 | 0 |
| Peptidoglycan polymer n subunits | cpd15665 | 0,03 |
| L Valine | cpd00156 | 0,35 |
| Isoheptadecanoylcardiolipin B subtilis | cpd15794 | 0,01 |

| | | |
|---|---|---|
| Zn2+ | cpd00034 | 0 |
| L-Glutamate | cpd00023 | 0,22 |
| L-Cysteine | cpd00084 | 0,08 |
| Bactoprenyl diphosphate | cpd02229 | 0,03 |
| dATP | cpd00115 | 0,02 |
| Fe2+ | cpd10515 | 0 |
| 10-Formyltetrahydrofolate | cpd00201 | 0 |
| L-Tryptophan | cpd00065 | 0,05 |
| GTP | cpd00038 | 0,14 |
| Ca2+ | cpd00063 | 0 |
| CoA | cpd00010 | 0 |
| Siroheme | cpd00557 | 0 |
| Mg+ | cpd00254 | 0 |
| Heme | cpd00028 | 0 |
| L-Proline | cpd00129 | 0,18 |
| NADP | cpd00006 | 0 |
| Spermidine | cpd00264 | 0 |
| FAD | cpd00015 | 0 |
| Glycine | cpd00033 | 0,51 |
| L-Tyrosine | cpd00069 | 0,12 |
| Stearoylcardiolipin B subtilis | cpd15793 | 0,01 |
| L-Glutamine | cpd00053 | 0,22 |
| L-Alanine | cpd00035 | 0,43 |
| L-Threonine | cpd00161 | 0,21 |
| NAD | cpd00003 | 0 |
| L-Leucine | cpd00107 | 0,38 |
| Cl- | cpd00099 | 0 |
| phosphatidylethanolamine dioctadecanoyl | cpd15533 | 0,01 |
| core oligosaccharide lipid A | cpd15432 | 0,03 |
| L-Serine | cpd00054 | 0,18 |
| RNA transcription | cpd17043 | 1 |
| ACP | cpd11493 | 0 |
| Dianteisoheptadecanoylphosphatidylglycerol | cpd15723 | 0,01 |
| L-Histidine | cpd00119 | 0,08 |
| Calomide | cpd00166 | 0 |
| Riboflavin | cpd00220 | 0 |
| Ubiquinone 8 | cpd15560 | 0 |
| L-Phenylalanine | cpd00066 | 0,15 |
| L-Lysine | cpd00039 | 0,29 |

**Supplementary Figure 1:** Genetic features captured by the model, classified according to COG categories

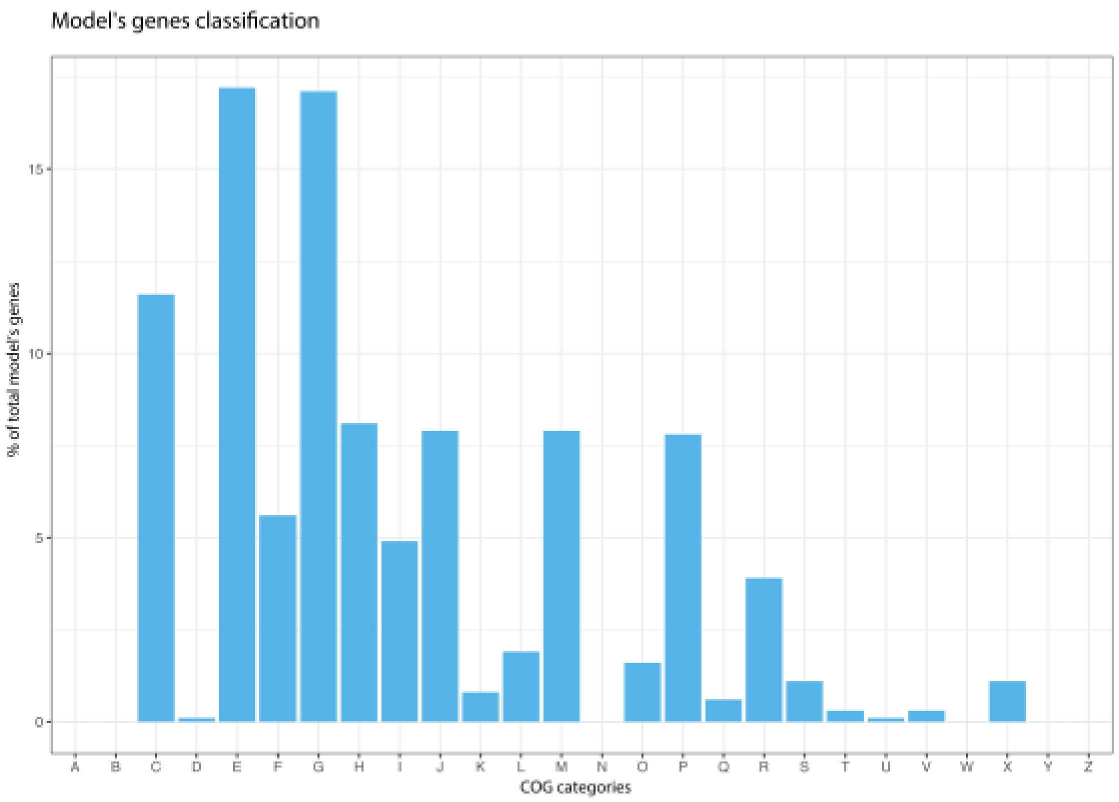


| | |
|---|---|
| A | RNA processing and modification |
| B | Chromatin structure and dynamics |
| C | Energy production and conversion |
| D | Cell cycle control, cell division, chromosome partitioning |
| E | Amino acid transport and metabolism |
| F | Nucleotide transport and metabolism |
| G | Carbohydrate transport and metabolism |
| H | Coenzyme transport and metabolism |
| I | Lipid transport and metabolism |
| J | Translation, ribosomal structure and biogenesis |
| K | Transcription |
| L | Replication, recombination and repair |
| M | Cell wall/membrane/envelope biogenesis |
| N | Cell motility |
| O | Posttranslational modification, protein turnover, chaperones |
| P | Inorganic ion transport and metabolism |
| Q | Secondary metabolites biosynthesis, transport and catabolism |
| R | General function prediction only |
| S | Function unknown |
| T | Signal transduction mechanisms |
| U | Intracellular trafficking, secretion, and vesicular transport |
| V | Defense mechanisms |
| W | Extracellular structures |
| X | No Functional Class Found |
| Y | Nuclear structure |
| Z | Cytoskeleton |

**Supplementary Figure 2:** Total amount of retained reactions per cut-off applied.

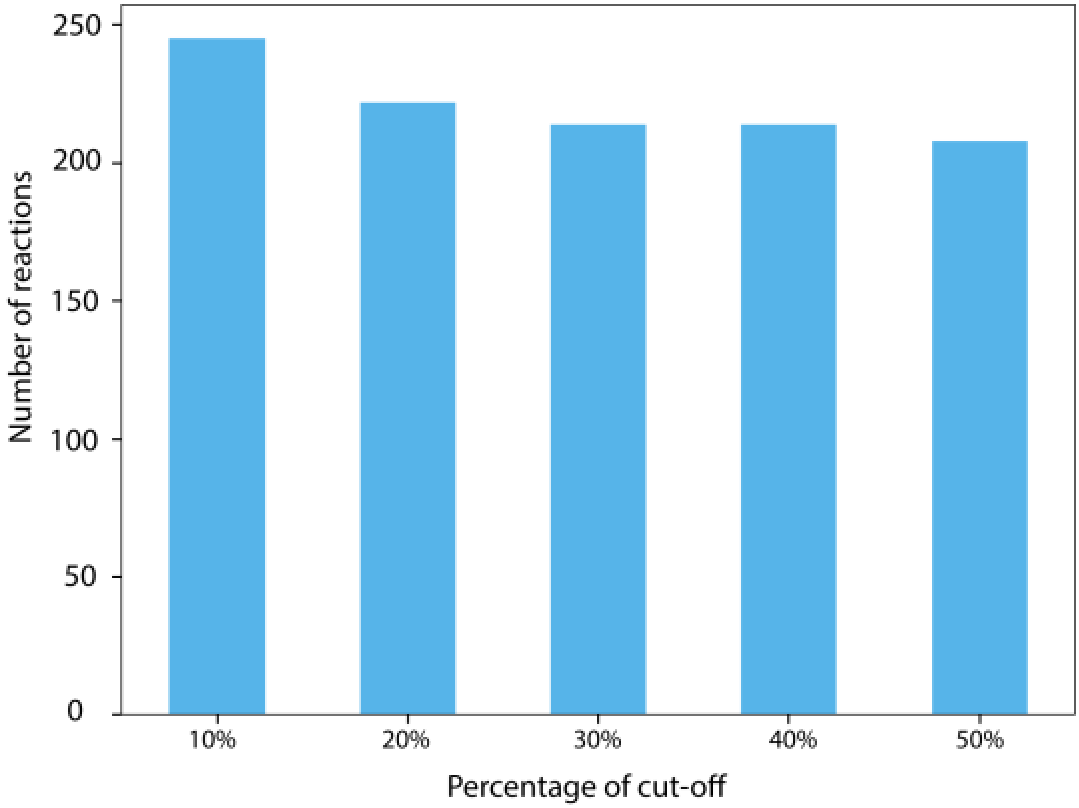

**Supplementary Figure 3:** Reactions increasing in flux (blue) and decreasing in flux (red) during the shift from soil to rhizosphere

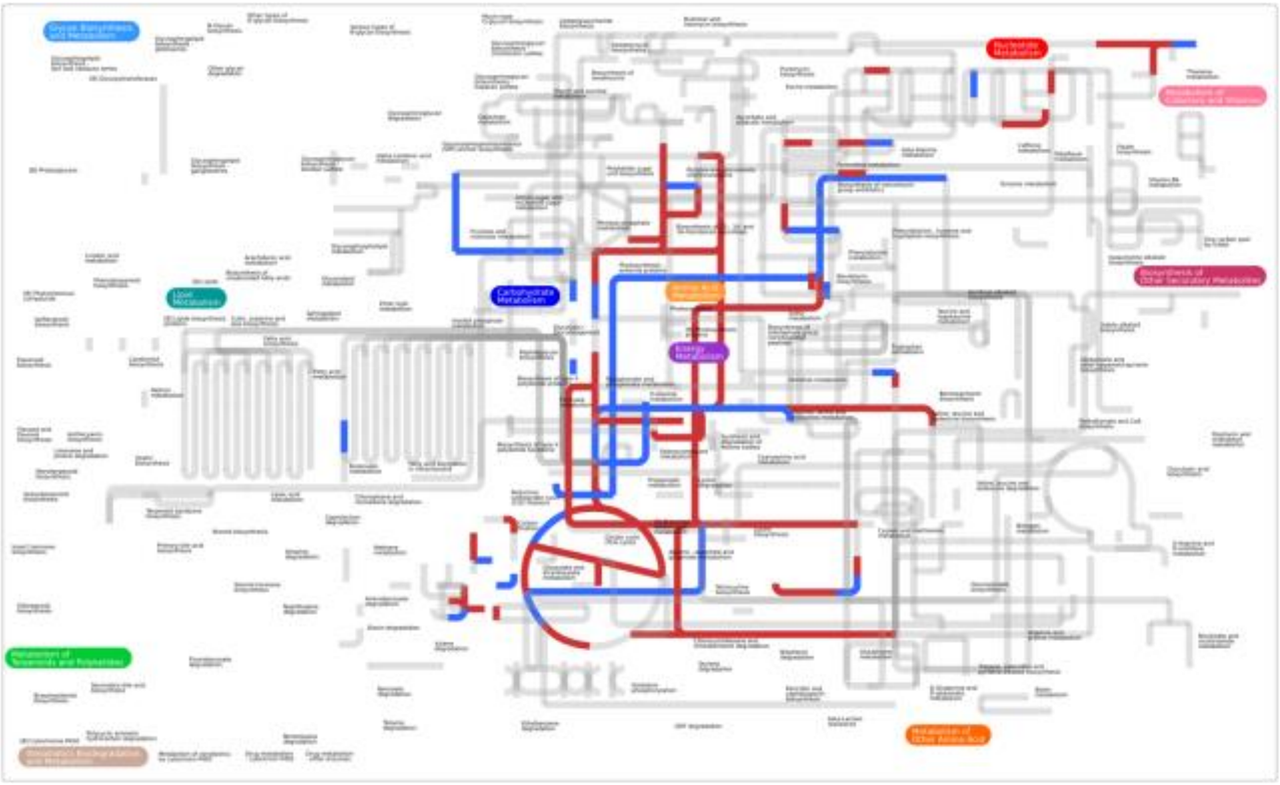
